# Adaptive co-transmission shapes sensory encoding and behavior

**DOI:** 10.64898/2026.10.12.758587

**Authors:** Andrea Cuentas-Condori, Sandeep Kumar, Peter Agoba, Patricia Chanabá López, Malcom Diaz Garcia, Daniel Colón-Ramos

## Abstract

Co-transmission expands the chemical repertoire of neurons, but whether coexisting neurotransmitters can be independently organized and remodeled by experience to control distinct aspects of behavior remains unclear. Here, we address this question in the AFD thermosensory neuron of *Caenorhabditis elegans*, which directs navigation toward a learned temperature preference. We find that glutamatergic and cholinergic vesicular transporters coexist within individual presynaptic boutons but differ in axonal trafficking, subsynaptic organization, and dependence on presynaptic machinery. Sensory experience differentially regulates the two systems: animals trained to prefer higher temperatures exhibit increased VGLUT/EAT-4 abundance, whereas those trained to prefer lower temperatures exhibit increased VAChT/UNC-17 abundance. The two transmitters also make distinct, context-dependent contributions to goal-directed navigation. Glutamatergic transmission promotes movement toward the learned thermal preference, whereas cholinergic transmission restrains continued migration beyond it. Analysis of locomotor trajectories further reveals that the two transmitters differentially regulate reorientation decisions across regions of the thermal gradient. Together, these findings identify *adaptive co-transmission*, a form of synaptic plasticity in which coexisting neurotransmitter systems are differentially organized and regulated by experience, and make distinct, context-dependent contributions to behavior.

## INTRODUCTION

Sensory representations are not fixed. Attention, motivation, arousal and prior experience can alter how sensory information is encoded and how animals respond to the same stimulus across contexts (Aton, 2013; McGann, 2015). Sensory systems must therefore integrate current sensory input with previous experience and behavioral goals to guide action. This flexibility can arise through changes in sensory transduction, neuronal excitability, synaptic strength and circuit connectivity, with experience-dependent plasticity occurring from peripheral sensory structures to central circuits (Akella et al., 2025; Clifford et al., 2007; Fu & Zuo, 2011; Tomko & Crapper, 1974). Among these mechanisms, changes in synaptic function provide a direct means of regulating how sensory information is transmitted and integrated within neural circuits. The synapse is therefore a critical site at which experience, behavioral state and chemical signaling converge to reshape information flow.

Neurotransmitter identity has traditionally been considered a defining and relatively stable feature of neuronal identity. This view was reinforced by a widely adopted interpretation of Dale’s principle (Dale, 1934), in which each neuron was thought to release a single neurotransmitter. Accordingly, neurons are commonly classified by their predominant transmitter (glutamatergic, cholinergic, GABAergic or monoaminergic) and that transmitter identity is considered relatively stable once established. However, growing evidence challenges both the notion that neurons are defined by a single neurotransmitter and the assumption that their transmitter identity remains fixed throughout life.

Many neurons across phyla co-express and release multiple small-molecule neurotransmitters (Granger et al., 2017; Trudeau & El Mestikawy., 2018; Vaaga et al., 2014). Moreover, transmitter expression can change in response to neuronal activity or experience (Bertuzzi et al., 2018; Dulcis et al., 2017; Dulcis & Spitzer, 2008; Li et al., 2024; Pratelli et al., 2024; Pratelli & Spitzer, 2025; Spitzer, 2015; Wu et al., 2020). These forms of flexibility in neurotransmitter usage can intersect. For example, experience can induce the addition or replacement of co-transmitters (Bertuzzi et al., 2018; Li et al., 2024), or alter their deployment across different axonal branches (Maddaloni et al., 2024). Importantly, coexisting transmitters can exhibit distinct release properties and forms of synaptic plasticity, and their contributions to synaptic signaling can be regulated (Hirai et al., 2024; Lee et al., 2010; Li et al., 2025; Ren et al., 2011; Shabel et al., 2014). Together, this growing body of work establishes neurotransmitter identity, spatial deployment, and functional contribution as potentially plastic features of neuronal signaling. A central unresolved question, however, is whether coexisting transmitter systems are independently organized and regulated within the same cellular and synaptic environment. In particular, it remains unknown whether experience can selectively remodel one transmitter system relative to another, and whether such transmitter-specific plasticity changes the information conveyed to downstream circuits and behavior.

To address these questions, we established an experimental system in which two endogenous transmitter systems can be examined within the same identified neuron and linked to an experience-dependent behavior. We focused on the AFD thermosensory neuron, which controls experience-dependent thermotaxis (Aoki & Mori, 2015; Goodman & Sengupta, 2018) and was identified in a broader cellular-resolution survey of neurotransmitter identity as a neuron with the molecular capacity for both glutamatergic and cholinergic transmission (Cuentas-Condori et al., 2025; Wang et al., 2024). The compact, genetically tractable nervous system of *C. elegans*, together with its mapped connectivity and quantitative behaviors, makes it possible to connect transmitter-specific cell biology within an identified neuron to circuit function and behavior. Using this system, we find that glutamatergic and cholinergic transmitter systems show distinct presynaptic organization, are differentially regulated by sensory experience and sensory transduction, and make context-dependent contributions to navigation. Together, these properties allow co-transmission to support flexible tracking of a learned sensory goal, a capacity that becomes particularly important when the sensory environment changes. Our findings identify adaptive co-transmission as a form of synaptic plasticity in which coexisting transmitter systems can be differentially organized and regulated to support experience-dependent behavior.

## RESULTS

### Glutamatergic and cholinergic machinery coexist within individual AFD presynaptic boutons

Our previous cellular-resolution survey of neurotransmitter identity unexpectedly identified both AFD neurons, AFDL and AFDR, among neurons in which endogenous *eat-4*/VGLUT and *unc-17*/VAChT reporters overlap (Cuentas-Condori et al., 2025). This observation motivated us to examine directly whether the molecular machinery for glutamatergic and cholinergic transmission coexists within identified AFD neurons. AFD is a bilateral pair of thermosensory neurons located in the head of *C. elegans* (Fig. 1A) that encodes experience-dependent animal navigation on a thermal gradient, the animal’s capacity to perform thermotaxis (Aoki & Mori, 2015; Goodman & Sengupta, 2018). Using an AFD-specific blue-fluorescent marker (AFD::BFP), we detected expression of endogenous reporters for *eat-4*/VGLUT, *unc-17*/VAChT and *cha-1*/ChAT in the same identified AFD neurons (Fig. 1B– D). Thus, AFD expresses the machinery required for glutamate vesicular loading together with both acetylcholine synthesis and vesicular loading.

**Figure 1.**
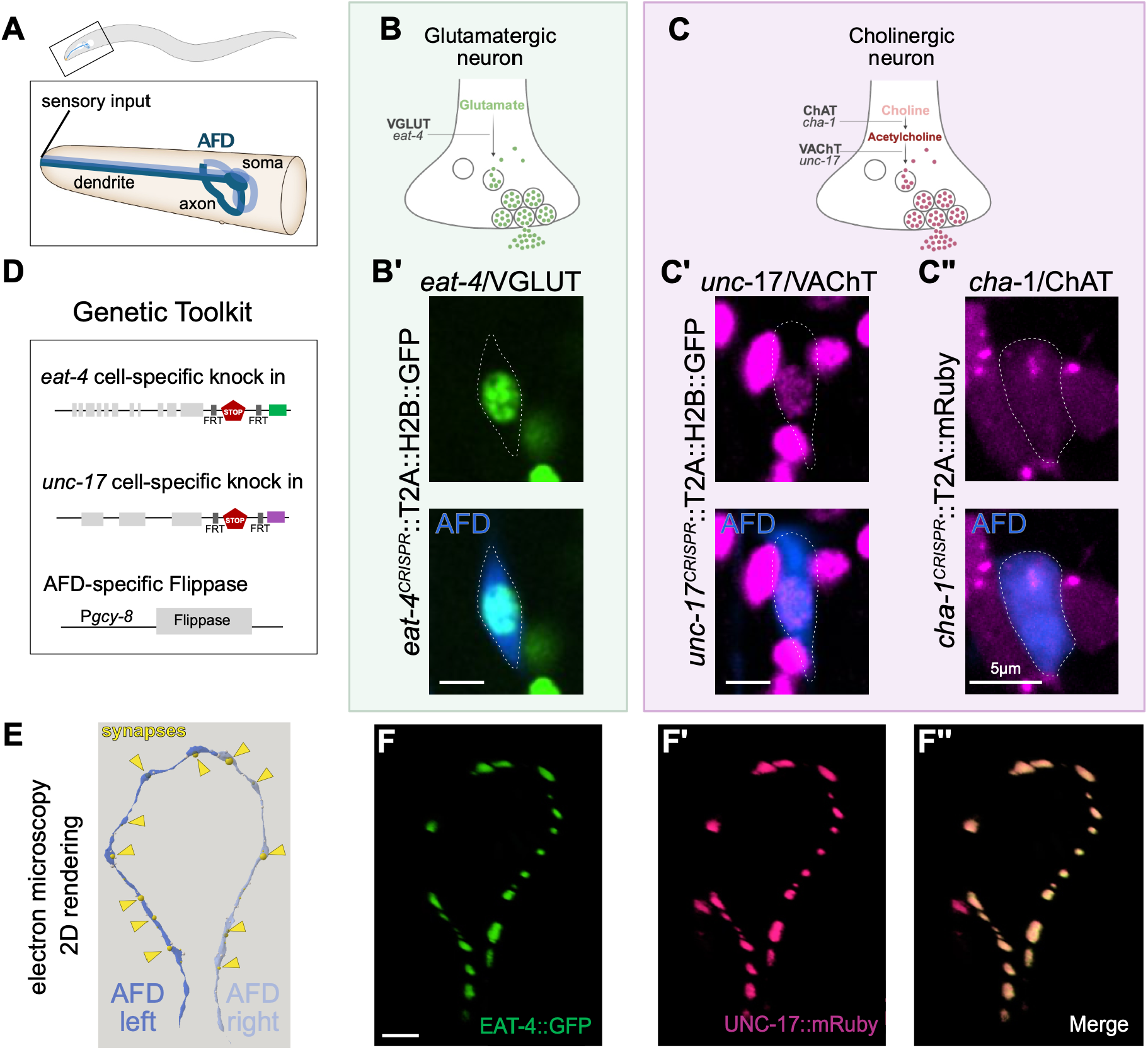
AFD neurons express the machinery to synthesize and release glutamate and acetylcholine. **(A)** AFD neurons (blue) localize in the *C. elegans* head and receive sensory input through their anterior ending. **(B)** Glutamatergic neurons are defined by the expression of the conserved Vesicular Glutamate Transporter (VGLUT/EAT-4). **(B’)** AFD neurons (blue) express an endogenous *eat-4*/VGLUT transcriptional reporter (green). Scale bar is 5 µm. **(C)** Cholinergic neurons are defined by the expression of the conserved Vesicular Acetylcholine Transporter (VAChT/UNC-17) and the acetylcholine synthesis enzyme Choline Acetyltransferase (ChAT/CHA-1). AFD neurons (blue) express an endogenous **(C’)** *unc-17*/VAChT and **(C’’)** *cha-1*/ChAT transcriptional reporter (magenta). Scale bar is 5 µm. **(D)** Genetic tools for conditionally labeling endogenous EAT-4/VGLUT with GFP and UNC-17/VAChT with mRuby in AFD neurons. Tools are from the SynaptoTagMe toolkit (Cuentas-Condori et al. 2026). **(E)** 2D-rendering of the AFD axons (purple) and synapses (yellow) from electron microscopy micrographs organizes in a ring-like structure. **(F)** 2D-rendering of a 3D projection of AFD axons. AFD-specific EAT-4::GFP (green) and UNC-17::mRuby (magenta) organize in a ring-like structure and occupy the same synaptic boutons. Scale bar is 5 µm.

These observations consolidate several independent lines of evidence from previous studies. AFD is an established glutamatergic neuron: EAT-4/VGLUT-dependent transmission from AFD regulates its postsynaptic partner AIY and thermotaxis behavior (Ohnishi et al., 2011). Transcriptomic studies subsequently detected *cha-1*/ChAT expression in AFD, both in the *C. elegans* single-cell atlas and through AFD-specific transcriptional profiling (Harris et al., 2023; Taylor et al., 2021). More recently, an endogenous *unc-17*/VAChT reporter was detected at low but reproducible levels in NeuroPAL-identified AFD neurons (Wang et al., 2024). Together with our direct detection of *eat-4, unc-17* and *cha-1* in AFD neurons, these findings establish that AFD contains the molecular machinery for both glutamatergic and cholinergic transmission.

We next asked whether both transmitter systems are deployed to the presynaptic compartment. Using cell-specific labeling strategies from the SynaptoTagMe toolkit (Cuentas-Condori et al., 2025), we independently visualized endogenous VGLUT/EAT-4 and VAChT/UNC-17 proteins with GFP in AFD (Fig. 1E; Supplementary Fig. 1A–B). Both transporters were readily detected in the AFD axon and concentrated in discrete presynaptic clusters. Their abundance, however, was asymmetric. Under matched imaging conditions, VGLUT/EAT-4::GFP fluorescence was consistently higher than VAChT/UNC-17::GFP fluorescence. This difference was apparent both in line scans along the AFD axon (Supplementary Fig. 1C) and when fluorescence was quantified within individual synaptic regions of interest (ROIs; Supplementary Fig. 1C’), yielding approximately five-fold and three-fold differences, respectively. Because line scans additionally sample regions between discrete synaptic clusters, subsequent comparisons of transporter abundance were performed within synaptic ROIs and averaged across boutons within each AFD axon.

Finally, to determine whether the two transporters are targeted to the same presynaptic sites, we simultaneously labeled endogenous VGLUT/EAT-4 with GFP and VAChT/UNC-17 with mRuby in AFD (Fig. 1E-F). Both transporters were distributed throughout the presynaptic region and were detected together at individual AFD boutons (Fig. 1E-F and Supplementary Fig. 1D-E). Thus, glutamatergic and cholinergic vesicular transport machinery is not segregated to different branches or subsets of AFD synapses but coexists within the same presynaptic boutons.

### VGLUT/EAT-4 and VAChT/UNC-17 show distinct subsynaptic organization and trafficking

Although VGLUT/EAT-4 and VAChT/UNC-17 are present within the same AFD synaptic boutons, their asymmetric synaptic levels raised the possibility that the two transporters are organized differently within individual presynaptic terminals. To resolve their subsynaptic organization *in vivo*, we adapted our conditional endogenous labeling tools for STED microscopy by tagging VGLUT/EAT-4 with SNAP Tag and VAChT/UNC-17 with Halo Tag, together with GFP-tagged Liprin-α/SYD-2, an active-zone scaffold used here as a fiducial marker of presynaptic release sites (Fig. 2A-B) (Zhen & Jin, 1999). Live STED imaging revealed non-identical distributions of VGLUT/EAT-4 and VAChT/UNC-17 within individual boutons (Fig. 2C). We therefore examined the relative position of each transporter across multiple boutons from individual animals (Fig. 2D). For each bouton, we identified the fluorescence maximum of Liprin-α/SYD-2, VGLUT/EAT-4 and VAChT/UNC-17 and measured the distance between each transporter maximum and the active-zone landmark (Fig. 2E). VGLUT/EAT-4 was consistently positioned closer to active-zone regions than VAChT/UNC-17, which displayed a broader and more distal distribution (Fig. 2F). Thus, even within the same presynaptic boutons, the two synaptic vesicle transporters systems occupy distinguishable subsynaptic domains. Our observations of distinct subcellular organization are consistent with the model in which distinct vesicular transporters are differentially regulated and might be localizing to different subpopulations of synaptic vesicles.

**Figure 2.**
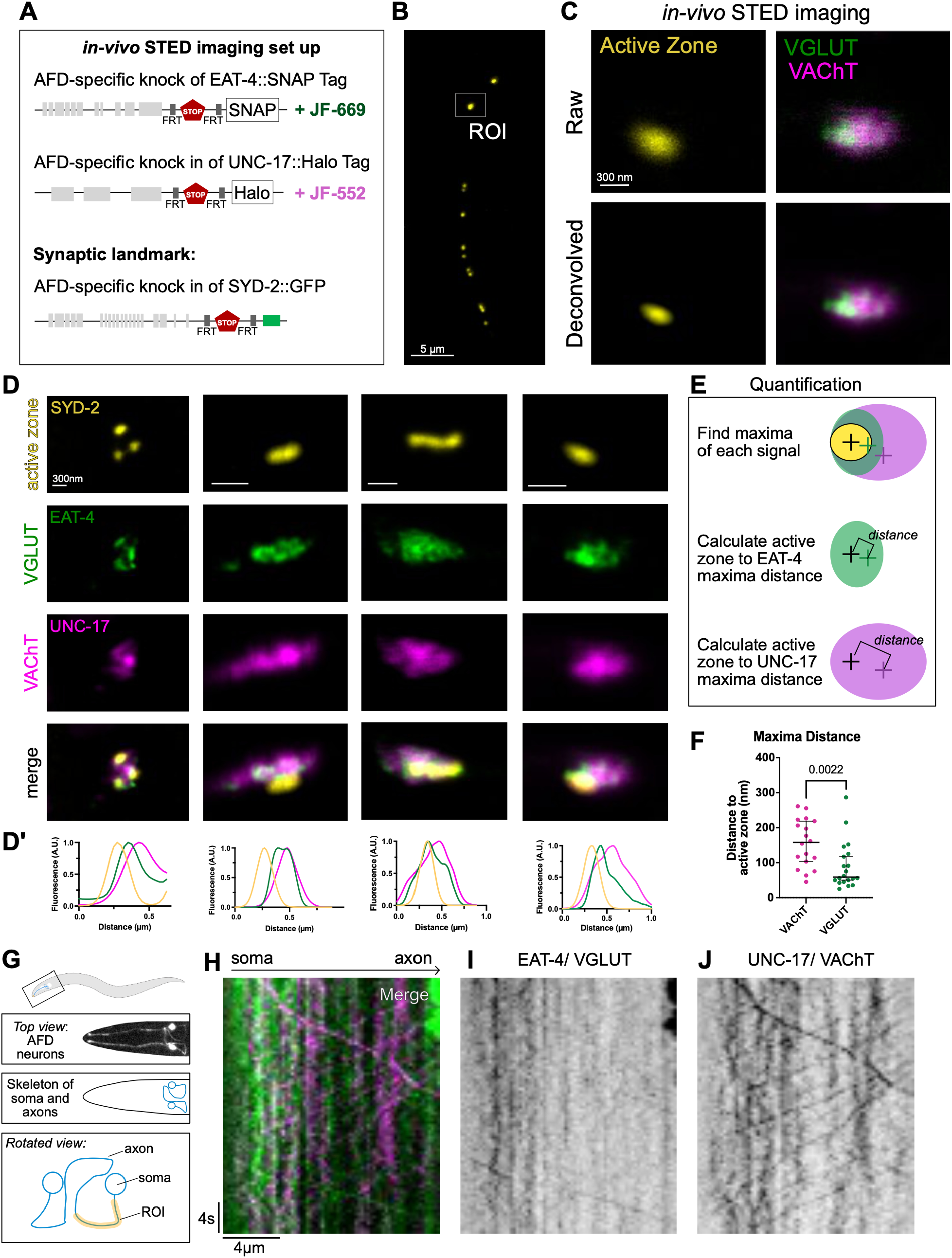
AFD neurons differentially organize EAT-4/VGLUT and UNC-17/VAChT. **(A)** Genetic tools for conditionally labeling of synaptic proteins compatible with STED microscopy. Endogenous EAT-4A/GLUT was labeled with SNAP-tag and UNC-17A/AChT with HaloTag specifically in AFD neurons. Janelia Fluor 669 and 552 dyes were used to label SNAP-tag and HaloTag, respectively. Endogenous SYD-2::GFP was used as an active zone landmark. (B) The endogenous active-zone marker SYD-2::GFP enables identification of individual boutons as regions of interest (ROIs) along the AFD axon. (C) Representative image of single bouton showing (top) raw data and (bottom) the corresponding image after deconvolution. Scale bar is 300 nm. (D) *In vivo* STED imaging of EAT-4A/GLUT (green), and UNC-17A/AChT (magenta), together with the active-zone landmark SYD-2. (D’) Corresponding line scans across individual boutons. Scale bar is 300 nm. (E) To quantify the position of each vesicular transporter relative to the active-zone landmark, fluorescence maxima were identified independently in each channel. (F) The distance between the fluorescence maxima of EAT-4A/GLUT or UNC-17A/AChT and the SYD-2 active-zone landmark differed significantly (p = 0.0022). (G) Top view of the AFD neurons and schematic representation of the AFD somas and axons. Bottom: magnified and rotated schematic showing the region of interest (ROI, yellow) used to monitor trafficking of synaptic proteins from the soma into the axon. (H) Representative merged kymograph of EAT-4A/GLUT and UNC-17A/AChT trafficking from the soma into the axon. Horizontal scale bar is 4um and vertical scale bar represents 4 seconds. (I) Kymograph of EAT-4 trafficking from the soma into the axon. (J) Kymograph of UNC-17 trafficking from the soma into the axon.

To further examine this regulation, we next asked whether this differential organization could arise before the transporters reach synapses. Using simultaneous two-color endogenous labeling, we followed VGLUT/EAT-4 and VAChT/UNC-17 trafficking from the AFD soma into the axon (Fig. 2G). Individual trafficking events for the two transporters were frequently non-coincident, producing distinct trajectories in two-color kymographs (Fig. 2H-J), consistent with at least partially independent transport from the cell body to the axon. The overall proportion of anterograde and retrograde events did not differ significantly between the two transporters (Supplementary Fig. 2A), and their anterograde and retrograde velocities were also similar (Supplementary Fig. 2B-C). These speed rates fall within the range reported for neuronal secretory cargo moving from the soma into *C. elegans* axons *in vivo* (Park *et al*., 2023). By contrast, both VGLUT/EAT-4 and VAChT/UNC-17 moved significantly more slowly than endogenous RAB3/RAB-3 particles within AFD (Supplementary Fig. 2B-C), further indicating that presynaptic material is not transported from the soma as a single uniform population.

Consistent with the possibility that transporter-specific sorting begins in the cell body, endogenous VGLUT/EAT-4 and VAChT/UNC-17 were enriched in neighboring but largely non-overlapping regions near the periphery of the AFD soma (Supplementary Fig. 2D). This observation is consistent with partial segregation of the two transporters before axonal delivery and raises the possibility that transmitter-specific organization begins during intracellular trafficking or biogenesis of the endo-vesicular structures in the soma.

We therefore asked whether the two transporters also depend differently on conserved components of the synaptic vesicle release machinery. Mutations in Syntaxin/*unc-64*, Munc13/*unc-13* and Munc18/*unc-18* produced distinct effects on synaptic VGLUT/EAT-4 and VAChT/UNC-17 abundance (Supplementary Fig. 2E-F). In Munc18/*unc-18* mutants, both transporters accumulated at synapses, whereas loss of Munc13/*unc-13* selectively increased VGLUT/EAT-4 without detectably altering VAChT/UNC-17. The selective dependence of VGLUT/EAT-4 on Munc13/*unc-13* is particularly notable given its preferential localization near Liprin-α/SYD-2-defined active zones. Together, these findings indicate that two vesicular neurotransmitter transporters sharing the same neuronal environment can nevertheless differ in their subcellular sorting, axonal trafficking, subsynaptic positioning and dependence on presynaptic release machinery. These differences are consistent with VGLUT/EAT-4 and VAChT/UNC-17 associating, at least in part, with distinct presynaptic transport or vesicle populations. Thus, the glutamatergic and cholinergic transmitter systems can be differentially organized and regulated within the same neuron.

### Sensory experience reconfigures transmitter-specific vesicular transporters on the timescale of behavioral adaptation

AFD is a central regulator of experience-dependent thermotaxis, in which animals navigate toward a learned temperature preference established by their prior cultivation (Aoki & Mori, 2015; Goodman & Sengupta, 2018; Hedgecock & Russell, 1975). We therefore asked whether sensory experience that establishes distinct thermotactic goals also regulates the presynaptic machinery of AFD’s two neurotransmitter systems. We first tested this idea by simultaneously monitoring endogenous VGLUT/EAT-4::GFP and VAChT/UNC-17::mRuby in AFD neurons of animals continuously cultivated at 15°C, 20°C, or 25°C (Fig. 3A). Sensory experience regulated the two vesicular transporters in opposite directions: VGLUT/EAT-4 abundance increased with warmer experiences (Fig. 3B–C), whereas VAChT/UNC-17 abundance decreased (Fig. 3D–E). Consequently, the relative abundance of the two vesicular transporters varied with the learned thermotactic goal, with VAChT/UNC-17 relatively enriched in animals trained to prefer lower temperatures and VGLUT/EAT-4 relatively enriched in animals trained to prefer higher temperatures. This relationship was preserved when the fluorescent tags were reciprocally exchanged, with VGLUT/EAT-4 labeled with mRuby and VAChT/UNC-17 with GFP (Supplementary Fig. 3A–B), confirming that the opposing regulation reflects the transporter systems rather than fluorophore identity.

**Figure 3.**
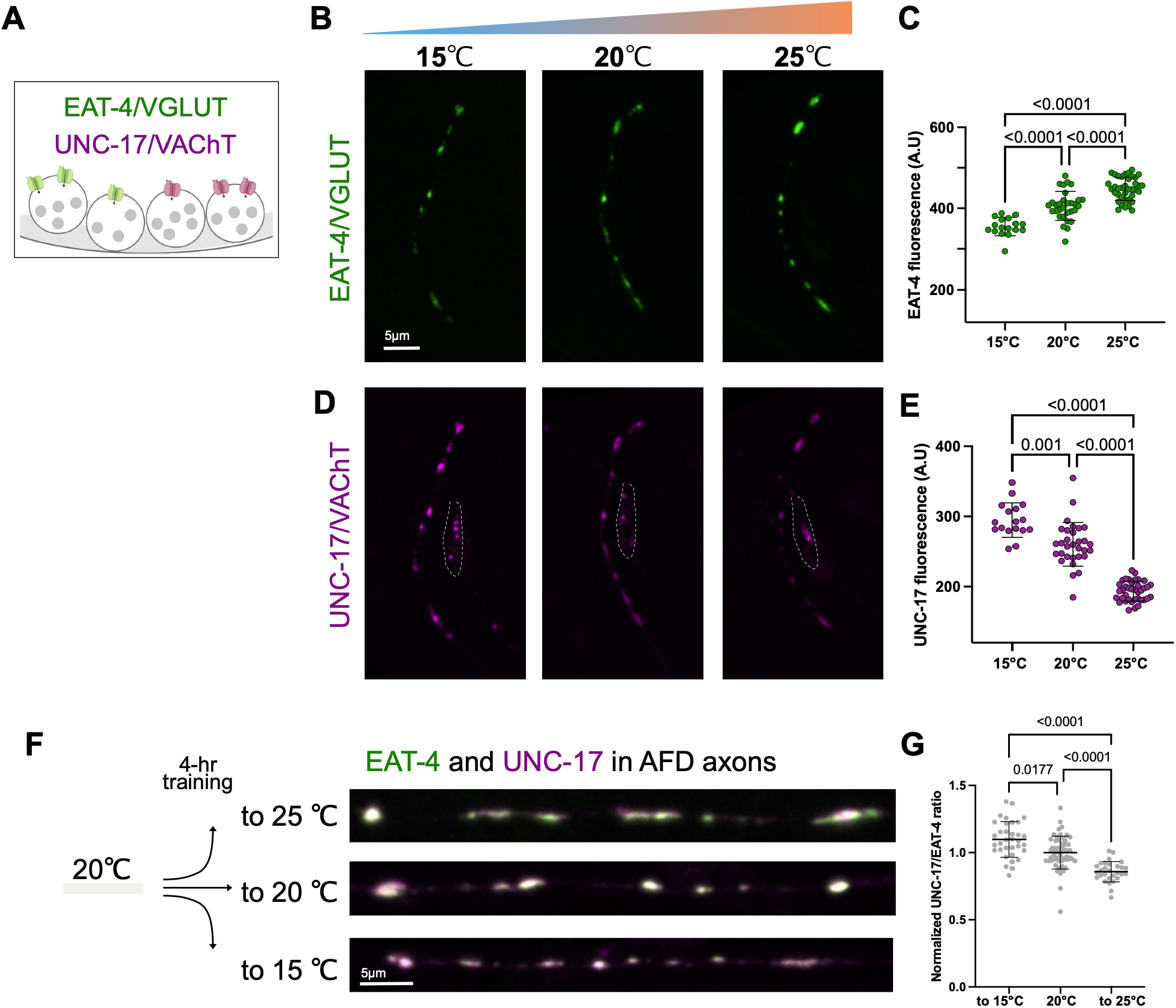
EAT-4/VGLUT and UNC-17/VAChT are differentially regulated by experience. **(A)** Dual labeling of EAT-4::GFP and UNC-17::mRuby in the thermosensory neuron AFD. **(B)** Warmer sensory experience upregulates detectable EAT-4/VGLUT levels in AFD axons. **(C)** Quantification of EAT-4::GFP across sensory experiences. Each dot represents an axon. **(D)** Warmer sensory experience downregulates UNC-17/VAChT levels in AFD axons. **(E)** Quantification of UNC-17::mRuby across sensory experiences. Each dot represents an axon. **(F)** Animals grown at 20°C and exposed for 4 hours to 15°C, 20°C, and 25°C show different UNC-17/EAT-4 ratios at synapses. **(G)** Quantification of UNC-17/EAT-4 ratios decreases with warmer experiences.

Previous work established that AFD synaptic output is shaped by thermal experience and contributes to thermotactic memory and learned preference. These studies showed experience-dependent changes in synaptic vesicle release (Samuel et al., 2003) and in functional coupling between AFD and its postsynaptic partner (Hawk et al., 2018), raising the possibility that changes in synaptic output could reflect a broader reconfiguration of presynaptic sites with experience. We therefore asked whether the opposing changes in VGLUT/EAT-4 and VAChT/UNC-17 (Fig. 3B-E) were accompanied by broader remodeling of AFD presynaptic components. Interestingly, the number of presynaptic active zones, measured using endogenous Clarinet/CLA-1, remained unchanged across thermal experiences (Supplementary Fig. 3C′). Endogenous RAB3/RAB-3, a synaptic-vesicle-associated protein, also did not show the opposing regulation observed for the two vesicular transporters, although its abundance decreased under the warmest condition (Supplementary Fig. 3C″). Moreover, exposure to the same cultivation temperatures did not alter VGLUT/EAT-4 abundance in the glutamatergic chemosensory neurons ASE or VAChT/UNC-17 abundance in the cholinergic chemosensory neuron ADF (Supplementary Fig. 3H-I). Thus, although AFD synaptic output is known to vary with thermal experience, the molecular remodeling observed here is not uniformly shared by the broader presynaptic components examined. Instead, thermal experience produces particularly pronounced and opposing regulation of the two coexisting transmitter systems specifically within AFD neurons.

We next asked whether these differences were fixed consequences of prolonged cultivation or whether the relative abundance of the two transmitter systems could adapt dynamically to a new sensory environment. Because experience-dependent thermotaxis, which is encoded through AFD activity (Mori & Ohshima, 1995), is remodeled over several hours after a change in cultivation temperature (Hedgecock & Russell, 1975) we asked whether the opposing regulation of VGLUT/EAT-4 and VAChT/UNC-17 develops within this behavioral adaptation window. Animals cultivated at 20°C were shifted as adults to 15°C, maintained at 20°C, or shifted to 25°C, and analyzed hourly (Supplementary Fig. 3D). Little or no change in the VAChT/UNC-17-to-VGLUT/EAT-4 ratio was detected during the first two hours, whereas differences began to emerge by approximately three hours and were robust after four hours of exposure to the new thermal environment (Supplementary Fig. 3E–E″; Fig. 3F-G). After 4 h, the VAChT/UNC-17-to-VGLUT/EAT-4 ratio increased following exposure to 15°C and decreased following exposure to 25°C (Fig. 3F–G). Thus, a few hours of new sensory experience are sufficient to alter the relative abundance of glutamatergic and cholinergic presynaptic machinery, on a timescale comparable to behavioral adaptation (Hedgecock & Russell, 1975).

We next asked whether this experience-dependent remodeling required synaptic activity through synaptic vesicle exocytosis. In mutants disrupting either *unc-18*/Munc18 or *unc-64*/Syntaxin, the increase in the UNC-17-to-EAT-4 ratio normally induced by a shift to 15 °C was no longer detected, whereas the decrease produced by a shift to 25 °C remained robust (Supplementary Fig. 3F–G). Thus, cold- and warm-dependent remodeling of the two transmitter systems are mechanistically separable. The requirement for intact exocytic machinery during cold adaptation is consistent with the possibility that ongoing use and recycling of transporter-containing synaptic vesicles contributes to remodeling their relative abundance at AFD synapses, whereas warm-dependent remodeling can proceed through an exocytosis-independent mechanism.

Together, these results identify a form of transmitter-specific presynaptic plasticity in which sensory experience reweights two coexisting neurotransmitter systems within the same neuron. Importantly, the mechanisms underlying this remodeling are themselves context dependent: cold-dependent reweighting requires intact exocytic machinery, whereas warm-dependent remodeling can proceed independently of it. Thus, co-transmitters can be regulated as partially independent units whose relative contribution is adjusted through distinct presynaptic mechanisms according to sensory experience. We refer to this capacity for context-dependent, transmitter-specific reconfiguration as adaptive co-transmission.

### AFD sensory transduction differentially regulates co-transmitter machinery

The previous experiments showed that thermal experience can reweight VGLUT/EAT-4 and VAChT/UNC-17 within AFD synapses (Fig. 3). We therefore asked whether this regulation depends on the sensory transduction pathway through which AFD detects temperature (Aoki & Mori, 2015; Goodman & Sengupta, 2018). In AFD, temperature-dependent changes in cGMP activate a cyclic nucleotide-gated channel composed of TAX-2 and TAX-4 subunits, coupling sensory input to calcium signaling and downstream cellular responses (Fig. 4A) (Kimura et al., 2004). We examined endogenous VGLUT/EAT-4 and VAChT/UNC-17 in animals lacking either component of this sensory transduction channel.

**Figure 4.**
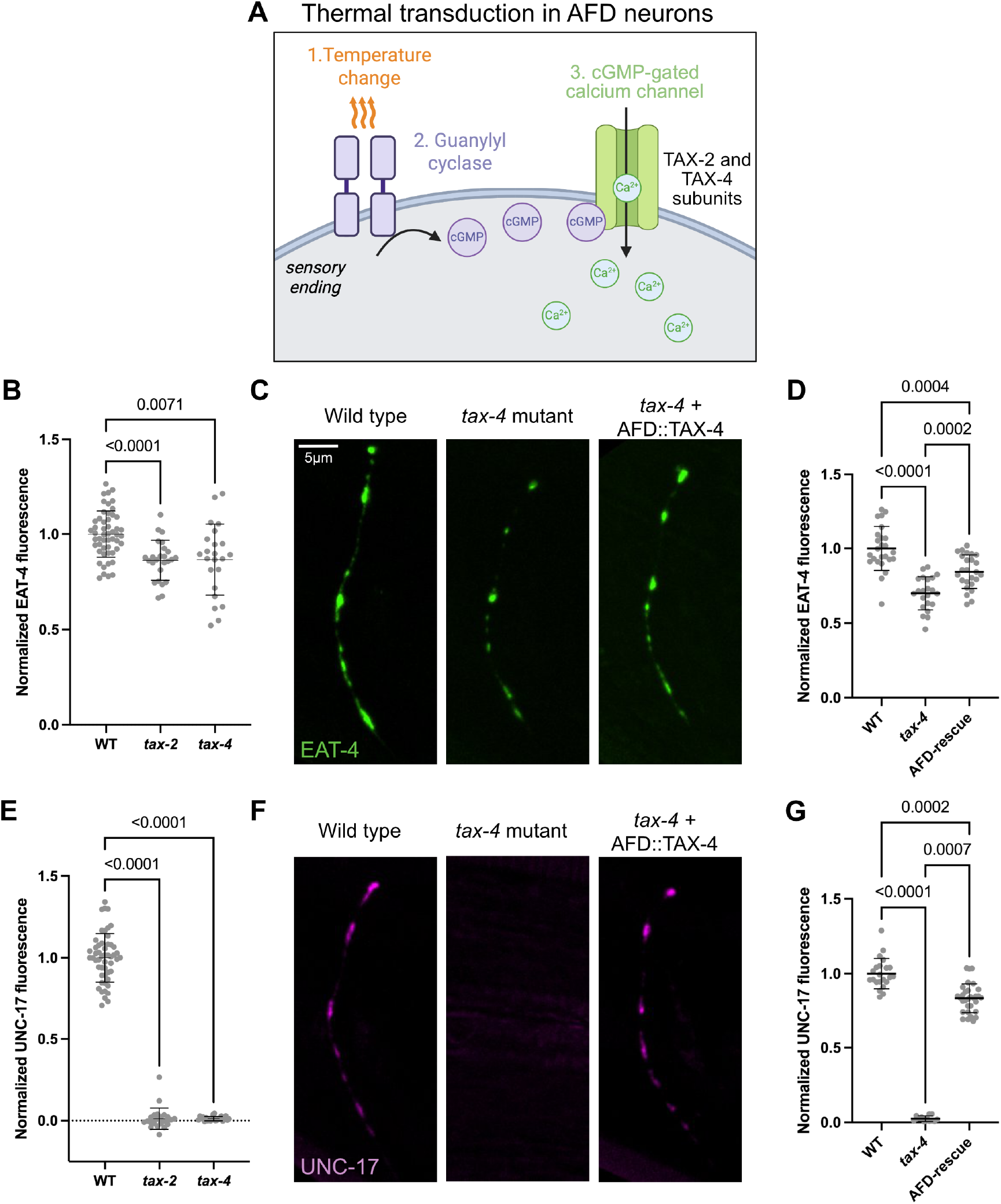
EAT-4 and UNC-17 are differentially and cell-autonomously regulated by sensory transduction. **(A)** Thermal transduction model in the AFD sensory ending. Temperature changes activate production of cGMP, which activates cGMP-activated ion channels encoded by *tax-4* and *tax-2* genes, required for sensory transduction. **(B)** Normalized EAT-4/VGLUT fluorescence in wild-type, *tax-2*, and *tax-4* mutant animals. **(C)** EAT-4::GFP fluorescence in AFD axons of (left) wild-type, (center) *tax-4* mutant, or (right) AFD-specific AFD::TAX-4 rescue. **(D)** Quantification of EAT-4/VGLUT levels in wild-type, *tax-4* mutants, and *tax-4*; AFD-specific TAX-4 rescue animals. **(E)** Normalized UNC-17/VAChT fluorescence in wild-type, *tax-2*, and *tax-4* mutant animals. **(F)** UNC-17::mRuby fluorescence in AFD axons of (left) wild-type, (center) *tax-4* mutant, or (right) AFD-specific AFD::TAX-4 rescue. **(G)** Quantification of UNC-17/VAChT levels in wild-type, *tax-4* mutants, and *tax-4*; AFD-specific TAX-4 rescue animals.

Loss of either *tax-2* or *tax-4* reduced synaptic VGLUT/EAT-4 levels in AFD (Fig. 4B). By contrast, the effect on the cholinergic transporter was substantially stronger: VAChT/UNC-17 became nearly undetectable in AFD axons of both mutants (Fig. 4E). To determine whether these effects arise from a requirement for sensory transduction within AFD itself, we restored TAX-4 specifically in AFD neurons. AFD-specific TAX-4 expression partially restored both VGLUT/EAT-4 and VAChT/UNC-17 levels (Fig. 4D-G), demonstrating a cell-autonomous requirement for sensory transduction in regulating both transmitter systems. Notably, however, the near-complete loss of VAChT/UNC-17 compared with the more moderate reduction in VGLUT/EAT-4 indicates that the two transmitter systems differ substantially in their dependence on sensory transduction. This differential sensitivity is consistent with the two transmitter systems being subject to distinct presynaptic regulatory mechanisms, in line with their differential organization and trafficking within AFD.

We next asked whether the loss of VAChT/UNC-17 reflected selective regulation of the vesicular transporter or a broader change in cholinergic identity. An endogenous transcriptional reporter for ChAT/*cha-1*, the enzyme required for acetylcholine synthesis, was strongly reduced in both *tax-2* and *tax-4* mutants (Supplementary Fig. 4B–B′). Thus, sensory transduction regulates multiple components of the cholinergic program in AFD rather than simply controlling VAChT/UNC-17 localization at synapses.

Loss of sensory transduction also alters broader components of AFD presynaptic architecture. We found that endogenous RAB3/RAB-3, a synaptic-vesicle-associated GTPase, was reduced in adult *tax-4* mutants (Supplementary Fig. 4A). This finding is consistent with previous reports that loss of TAX-2/TAX-4 signaling alters RAB-3 organization in AFD synapses (Hellman and Shen, 2011). Together, these observations support the idea that sensory transduction contributes to the organization of the broader synaptic-vesicle compartment in AFD, in addition to its differential effects on VGLUT/EAT-4 and VAChT/UNC-17.

We similarly observed changes in active-zone organization. In adult *tax-4* mutants, levels of endogenous Clarinet/CLA-1, a conserved active-zone scaffold (Xuan et al., 2017), were reduced while the number of detectable CLA-1 puncta increased (Supplementary Fig. 4E–F). This phenotype closely parallels previous observations for another conserved active-zone protein, Liprin-α/SYD-2 (Zhen & Jin, 1999), which was also reported to form more numerous but less intense puncta in *tax-2* and *tax-4* mutant AFD neurons (Hellman and Shen, 2011). Together, these findings support the idea that TAX-2/TAX-4-dependent sensory transduction contributes to the organization of active-zone components at AFD synapses.

Developmental analysis further distinguishes these broader effects on presynaptic architecture from the regulation of individual transmitter systems. Hellman and Shen found that excess SYD-2 puncta were already present throughout larval development in *tax-2* and *tax-4* mutants, whereas an increase in RAB-3 puncta emerged later, beginning around the L3 stage. Consistent with an early effect on active-zone organization, we found that first-stage (L1) *tax-4* larvae already displayed reduced CLA-1 abundance and increased numbers of CLA-1 puncta (Supplementary Fig. 4E′–F′). Strikingly, however, the two vesicular neurotransmitter transporters behaved differently at this same developmental stage: VAChT/UNC-17 was already strongly depleted, whereas VGLUT/EAT-4 levels were indistinguishable from wild type (Supplementary Fig. 4C′–D′). In adults, both transporters were reduced, although VAChT/UNC-17 was largely undetectable (Supplementary Fig. 4C–D). Thus, the differential effects on VGLUT/EAT-4 and VAChT/UNC-17 cannot be explained simply by a uniform defect in presynaptic assembly.

Together, these results extend previous evidence that TAX-2/TAX-4 signaling is required for normal presynaptic architecture in AFD by revealing an additional level of specificity: the two neurotransmitter systems occupying these synapses have distinct dependencies on the same sensory transduction pathway. Cholinergic machinery shows a particularly strong and early requirement for TAX-2/TAX-4 signaling, whereas the dependence of VGLUT/EAT-4 becomes more pronounced later in development. Thus, sensory transduction regulates not only the organization of the presynaptic terminal, but also the relative availability of the chemical signaling systems operating within it.

### Co-transmitters regulate goal-directed navigation in distinct sensory contexts

AFD contributes to both positive and negative thermotaxis (Aoki & Mori, 2015; Goodman & Sengupta, 2018), but transmitter-specific studies have directly linked its glutamatergic output to cold-directed navigation. VGLUT/EAT-4-dependent glutamate release from AFD inhibits the postsynaptic AIY interneuron and biases animals toward colder temperatures, providing a defined behavioral context in which to examine how a second transmitter from the same neuron contributes to navigation (Ohnishi et al., 2011). We therefore focused on negative thermotaxis. Animals adapted to 15 °C and subsequently placed near 20°C on a 15–25 °C thermal gradient migrate down the gradient, toward their learned thermal goal (Fig. 5A–A′). AFD sensory responses are themselves shaped by experience, with gain preferentially enhanced within behaviorally relevant regions of the thermal landscape (García et al., 2026). We therefore asked whether glutamate and acetylcholine make distinct contributions as animals move through different sensory contexts during goal-directed navigation.

**Figure 5.**
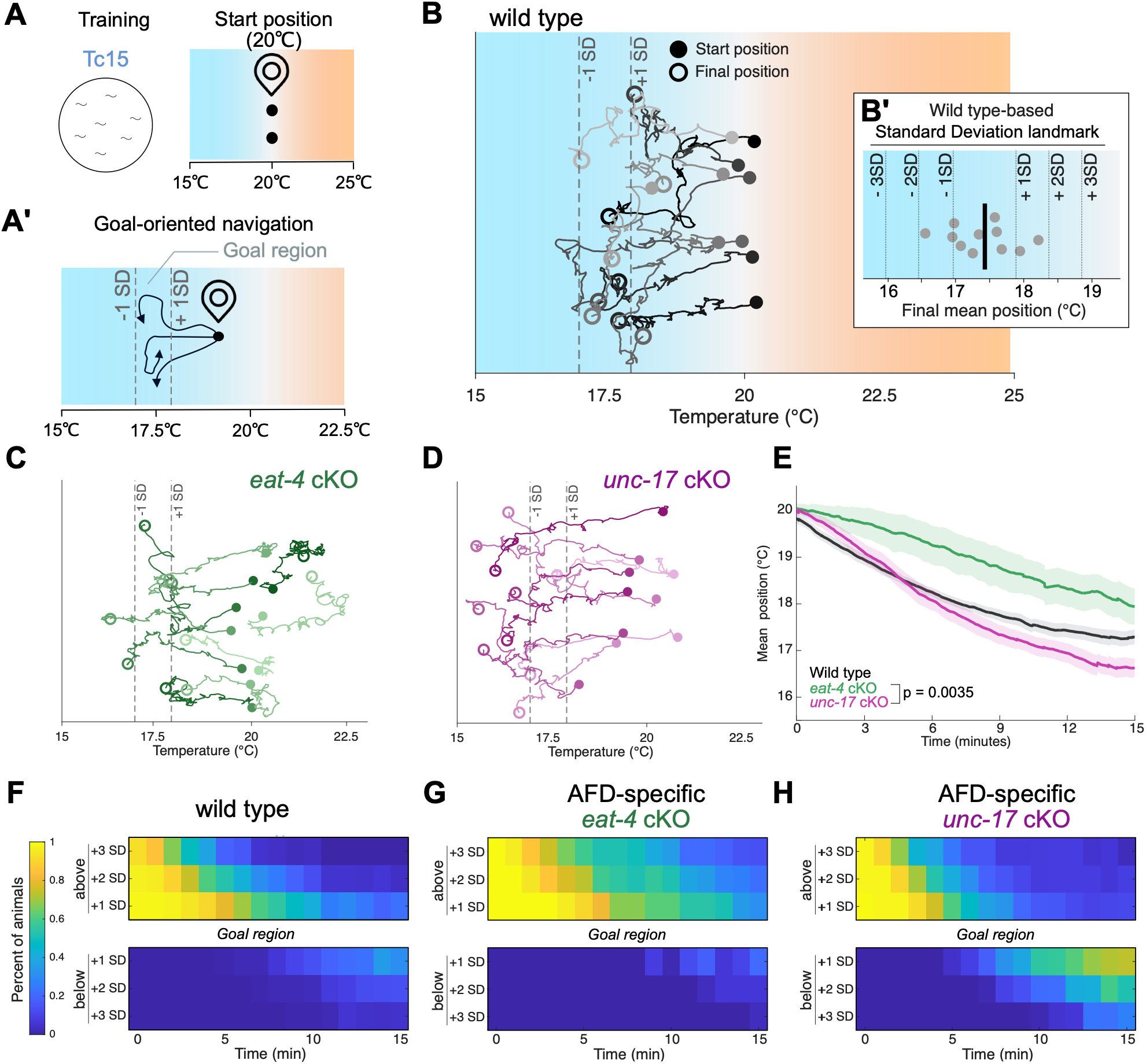
EAT-4/VGLUT and UNC-17A/AChT differentially regulate goal-oriented navigation. **(A)** Animals cultivated at 15°C (Tc15) were tested on a linear 15-25°C thermal gradient, with all animals starting at 20°C. (A’) Tc15 animals perform negative thermotaxis, migrating toward colder temperatures and their learned thermal preference. (B) Representative trajectories often wild-type animals during a 15-min thermotaxis assay after Tc15 training. Filled circles indicate starting positions and open circles indicate final positions. The -1 SD and +1 SD landmarks were defined from the distribution of mean thermal positions across eleven independent wild-type plates during the final 5 min of the assay. (B^1^) Mean thermal position of the eleven wild-type plates used to define these landmarks. **(C)** Representative trajectories often AFD-specific *eat-4cKO* mutant animals during a 15-minute thermotaxis assay after Tc15 training. The -1 SD and +1 SD landmarks correspond to the wild-type-derived landmarks defined in B-B’. (D) Representative trajectories of ten AFD-specific *unc-17cKO* mutant animals during a 15-minute thermotaxis assay afterTc15 training. The -1 SD and +1 SD landmarks correspond to the wild-type-derived landmarks defined in B-B’. (E) Mean thermal position of wild-type (black), AFD-specific *eat-4* cKO (green), and AFD-specific *unc-17* cKO (magenta) animals over the course of the assay. **(F)** Heatmaps showing the percentage of wild-type animals located (top) on the warmer side of and (bottom) on the colder side of the wild-type-derived landmarks defined in B-B’. (G) Heatmaps showing the percentage of AFD-specific *eat-4* cKO animals located (top) on the warmer side of and (bottom) on the colder side of the wild-type-derived landmarks defined in B-B’. **(H)** Heatmaps showing the percentage of AFD-specific *unc-17* cKO animals located (top) on the warmer side of and (bottom) on the colder side of the wild-type-derived landmarks defined in B-B’.

Wild-type animals followed stereotyped trajectories from their starting position near 20°C and progressively accumulated within a restricted region around 17–18 °C after 15 minutes (Fig. 5B–), a time window previously shown to capture goal-directed progression toward learned thermal preferences and changes in navigation as animals approach their thermal goal (García et al., 2026; Luo et al., 2014).

Animals lacking VGLUT/EAT-4 specifically in AFD remained displaced toward warmer temperatures and approached the wild-type learned goal region more slowly (Fig. 5C, 5E; Supplementary Fig. 5A–C). This phenotype is consistent with previous work showing that loss of glutamatergic transmission from AFD biases thermotaxis toward warmer temperatures (Ohnishi et al., 2011) and extends those findings by showing that AFD glutamate is particularly important for efficient progression from the warm starting region toward the learned thermal goal. By contrast, animals lacking VAChT/UNC-17 specifically in AFD initially migrated down the gradient similarly to wild type but failed to remain within the learned goal region where wild-type animals accumulated, instead continuing farther down the gradient and becoming progressively enriched at temperatures below the learned preference (Fig. 5D–E). These results identify acetylcholine as a second AFD output required not simply for cold-directed movement, but for appropriately constraining that movement as animals approach and pass their learned thermal goal. Thus, glutamate and acetylcholine make complementary, context-dependent contributions to negative thermotaxis: glutamate promotes effective navigation towards the goal region, whereas acetylcholine helps prevent continued migration beyond the goal region. Together, these findings suggest that successful goal-directed navigation emerges from the coordinated action of two transmitter systems within the same sensory neuron, with each becoming particularly important in different regions of the sensory landscape.

To resolve when and where these navigation defects emerged, we used the wild-type distribution assay (Fig. 6B’) to define temperature landmarks around the wild-type goal region and followed, minute by minute, the fraction of each population crossing these landmarks (Fig. 5B′, F–H). On the warm side of the gradient, wild-type and AFD-specific *unc-17* cKO animals efficiently progressed towards the learned goal region, whereas *eat-4* cKO animals crossed them more slowly and a fraction remained in gradient areas before the goal region throughout the assay (Fig. 5F–H; Supplementary Fig. 5D–F). Conversely, after reaching the goal region, *unc-17* cKO animals increasingly crossed landmarks below the wild-type goal region and remained enriched at lower temperatures (Fig. 5H; Supplementary Fig. 5G–I). Thus, the two transmitter-dependent phenotypes are spatially distinct. These spatially distinct phenotypes correspond to different demands of goal-directed navigation: glutamatergic transmission is particularly important as animals progress toward their learned goal, whereas cholinergic transmission becomes important as they approach that goal and must limit continued movement beyond it.

**Figure 6.**
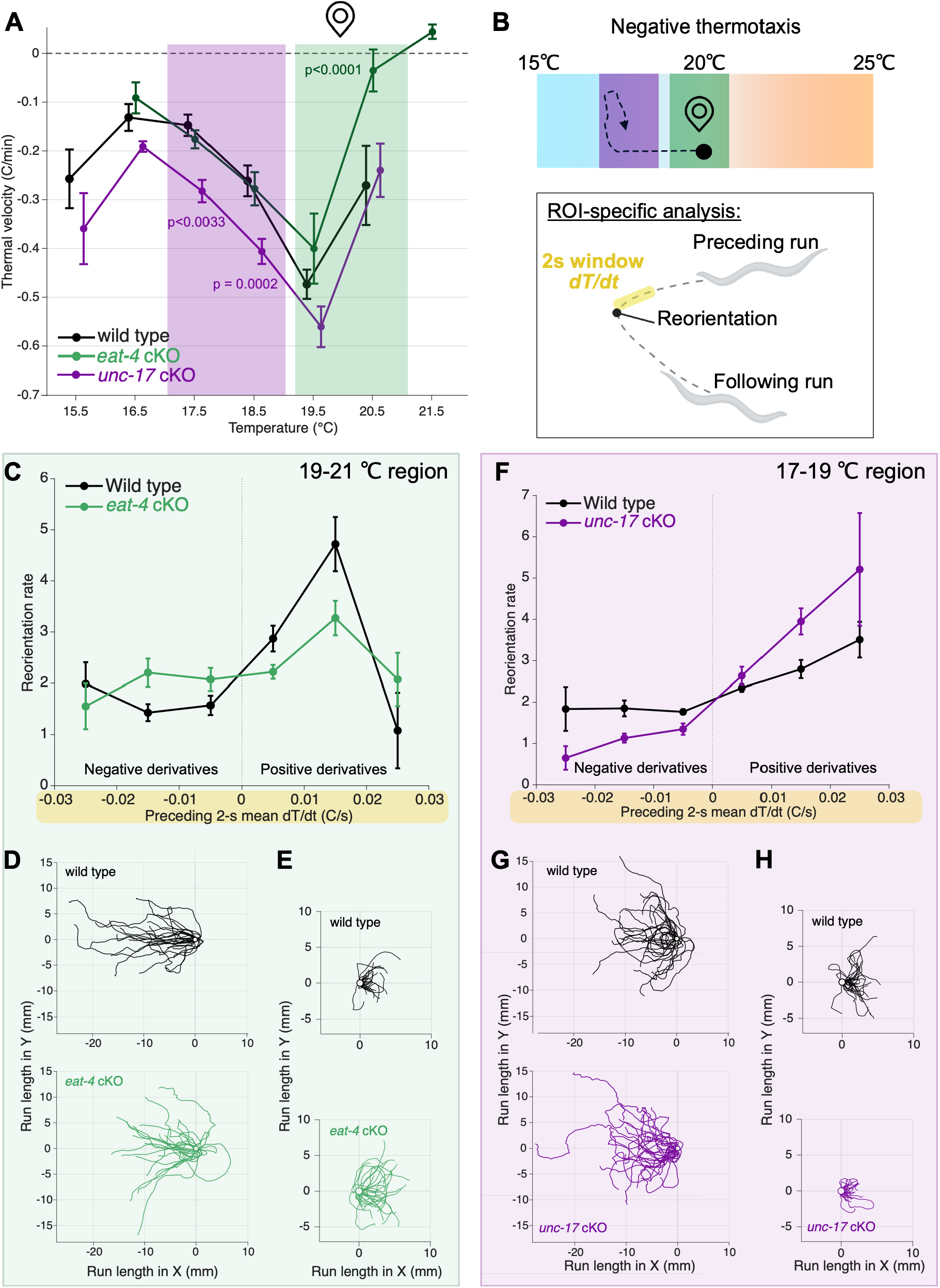
EAT-4/VGLUT and UNC-17/VAChT regulate context-specific features of goal-oriented navigation. **(A)** Thermal velocity across temperature bins for wild-type (black), AFD-specific *eat-4* cKO (green), and AFD-specific *unc-17* cKO (magenta) animals. The location-marker icon indicates the approximate starting position of animals in the assay. The genotype * temperature interaction across all three genotypes was tested using a hierarchical Gaussian linear mixed-effects model and likelihood-ratio test (p<0.0001). Within-temperature-bin comparisons between wild type and each mutant were obtained from hierarchical mixed-effects model contrasts with Satterthwaite-adjusted degrees of freedom and Holm correction for multiple comparisons across eligible temperature bins. Error bars represent SEM across independent assay plates. (B) Model and analysis framework for context-specific contributions of AFD glutamate and acetylcholine during negative thermotaxis. Animals begin near 20 °C and navigate down the thermal gradient toward their learned thermal goal. Glutamatergic transmission is particularly important within the warmer 19-21 °C region, whereas cholinergic transmission is particularly important within the cooler 17-19 °C region. For each reorientation, the mean thermal derivative (dT/dt) experienced during the preceding 2 s was calculated from the preceding run and related to reorientation rate and the direction of the following run. (C) Reorientation rate as a function of the mean dT/dt experienced during the preceding 2 seconds for wild type (black) and AFD-specific *eat-4* cKO (green) animals within the 19-21C region. Negative dT/dt values indicate movement associated with decreasing temperature, whereas positive values indicate movement associated with increasing temperature. Reorientation counts were analyzed using a Poisson generalized linear mixed-effects model with genotype, dT/dt bin and their interaction as fixed effects, exposure time as a log offset, and plate as a random intercept. The genotype x dT/dt-bin interaction was evaluated using a joint Wald test (p < 0.0001). (D) Thirty longest runs down the thermal gradient for wild-type (black) and AFD-specific *eat-4* cKO (green) animals within the 19-21 °C region. All tracks were translated to a common origin. (E) Thirty longest runs up the thermal gradient for wild-type (black) and AFD-specific *eat-4* cKO (green) animals within the 19-21 °C region. All tracks were translated to a common origin. (F) Reorientation rate as a function of the mean dT/dt experienced during the preceding 2 s for wild-type (black) and AFD-specific *unc-17* cKO (magenta) animals within the 17-19 °C region. Negative and positive dT/dt values indicate decreasing and increasing temperature, respectively. Statistical analysis was performed as in C. The genotype x dT/dt-bin interaction was evaluated using a joint Wald test (p < 0.0001). (G) Thirty longest runs down the thermal gradient for wild-type (black) and AFD-specific *unc-17* cKO (magenta) animals within the 17-19 °C region. All tracks were translated to a common origin. (H) Thirty longest runs up the thermal gradient for wild-type (black) and AFD-specific *unc-17* cKO (magenta) animals within the 17-19 °C region. All tracks were translated to a common origin.

### Glutamate and acetylcholine bias distinct navigation strategies across the thermal landscape

Together, the trajectory and landmark analyses revealed a spatial organization to the two transmitter-dependent phenotypes (Fig. 5): loss of AFD glutamate most strongly disrupted navigation in warmer portions of the gradient, whereas loss of AFD acetylcholine produced its strongest defect at cooler temperatures. We therefore asked which components of navigation are differentially affected by each transmitter across these distinct thermal contexts. We first quantified thermal velocity (°C/min) across the gradient to determine how the effect of each transmitter varied with temperature. This analysis revealed a significant genotype-by-temperature interaction, with the *eat-4* cKO phenotype most pronounced at warmer temperatures and the *unc-17* cKO phenotype emerging more strongly at cooler temperatures (Fig. 6A). Guided by this spatial dependence, we examined navigation dynamics within a warmer 19–21 °C window and a cooler 17–19 °C window (Fig. 6B).

We next asked how recent thermal experience is translated into reorientation decisions. AFD is well established to encode temporal changes in temperature. In freely moving animals, short-term adaptation makes AFD activity sensitive to the temporal derivative of temperature, providing information about whether an animal is moving up or down a thermal gradient that could be used to bias reorientation behavior (Clark et al., 2007). Subsequent work showed that AFD integrates temperature changes over behaviorally relevant timescales comparable to those of turning decisions (Tsukada et al., 2016), and more recent measurements indicate that AFD acts as a leaky integrator of recent temperature changes whose gain is enhanced near the learned thermal preference (Díaz García et al., 2026). These observations motivated us to ask whether glutamate and acetylcholine differentially couple recent temperature change to reorientation behavior. We therefore quantified reorientation rate as a function of the mean dT/dt experienced during the 2 s immediately preceding each reorientation (Fig. 6B).

The two transmitters affected this sensory-to-behavior relationship in opposite ways. In the warmer 19–21 °C region, wild-type animals reoriented relatively infrequently following negative dT/dt, when temperature was decreasing, and more frequently following positive dT/dt, when temperature was increasing. Removing VGLUT/EAT-4 weakened this distinction: *eat-4* cKO animals reoriented more frequently than wild type following negative dT/dt and less strongly in response to positive dT/dt (Fig. 6C and Supplementary Fig. 6A). Thus, glutamatergic transmission reinforces a behavioral bias that sustains cooling trajectories while interrupting trajectories associated with increasing temperature, which is captured in the increased capacity of *eat-4* cKO animals to terminate runs down the gradient compared to wild type and *unc-17* cKO animals (Fig. 6D-E and Supplementary Fig. 6B).

In the cooler 17–19 °C region, loss of VAChT/UNC-17 shifted this relationship in the opposite direction. *unc-17* cKO animals reoriented less frequently than wild type following negative dT/dt and more frequently following positive dT/dt (Fig. 6F; and Supplementary Fig. 6C). Consequently, cooling trajectories were preferentially maintained, whereas trajectories associated with increasing temperature were more readily interrupted (Fig. 6G-H; and Supplementary Fig. 6D).

Thus, the two co-transmitters oppositely regulate how AFD’s derivative-based sensory representation is translated into behavioral decisions: glutamate promotes continued negative thermotaxis in warmer conditions, whereas acetylcholine restrains this behavior closer to the learned goal. These opposing effects suggest that AFD co-transmission may keep navigation referenced to a sensory goal, rather than simply reinforcing movement in a fixed direction. We therefore tested this model by changing the spatial relationship between the animal and its learned thermal goal.

### AFD co-transmitters support adaptive tracking of a learned thermal goal

To test whether the opposing actions of glutamate and acetylcholine are critical to maintain navigation relative to a learned goal, we changed the location of the goal while keeping the learned temperature unchanged. Animals trained at 15 °C were allowed to navigate for 15 min on a 15–25 °C gradient, after which the gradient was switched while the learned temperature remained unchanged (Fig. 7A). This manipulation displaced the learned goal to the opposite side of the arena, requiring animals to update the direction of navigation without changing the goal itself. Before the switch, *eat-4* cKO animals remained warmer than wild type, whereas *unc-17* cKO animals progressed farther down the gradient toward colder temperatures (Fig. 7B; Supplementary Fig. 7A). Following gradient reversal, wild-type animals redirected their movement toward the relocated goal, whereas both transmitter-specific mutants failed to appropriately update their navigation (Fig. 7B–D). Thus, both glutamatergic and cholinergic transmission from AFD are required for successful behavioral adjustment to a changing sensory environment.

**Figure 7.**
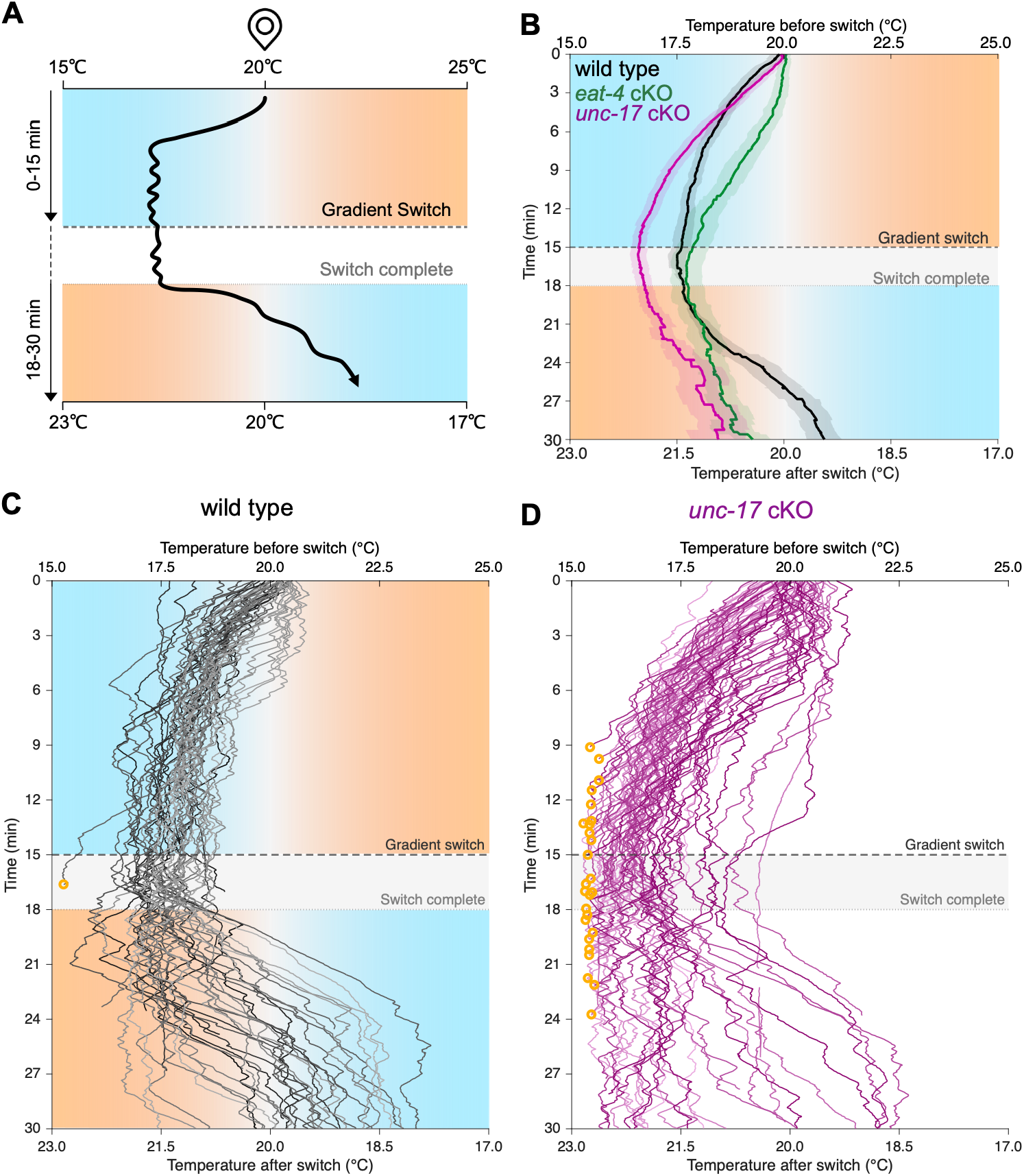
AFD co-transmitters coordinate adaptive tracking of a learned thermal goal. **(A)** Schematic of the gradient-switch assay. Animals trained at 15 °C were placed near 20 °C on a linear 15–25 °C thermal gradient and allowed to perform negative thermotaxis for 15 min. The direction of the thermal gradient was then reversed while maintaining 20 °C near the center of the arena. Gradient reversal was initiated at 15 min and completed by 18 min, generating a 23–17 °C gradient of opposite orientation. Animals were subsequently tracked until 30 min. **(B)** Mean thermal position over time for wild-type (black), AFD-specific *eat-4* cKO (green), and AFD-specific *unc-17* cKO (magenta) animals before and after gradient reversal. The upper x-axis indicates temperature before the switch and the lower x-axis indicates temperature after the switch. Dashed and dotted horizontal lines indicate initiation and completion of gradient reversal, respectively. Shaded regions represent SEM. **(C)** Individual thermal trajectories of wild-type animals during the gradient-switch assay. Each line represents one animal; orange circles indicate trajectories that exited the recording arena. Temperatures corresponding to animal position before and after gradient reversal are indicated on the upper and lower x-axes, respectively. *n* = 60 tracks from 5 independent assay plates. **(D)** Individual thermal trajectories of AFD-specific *unc-17* cKO animals during the gradient-switch assay, plotted as in C. Orange circles indicate trajectories that exited the recording arena. *n* = 82 tracks from 7 independent assay plates.

The cholinergic phenotype was particularly informative. Individual trajectories revealed that AFD-specific *unc-17* cKO animals frequently progressed beyond the region occupied by wild-type animals before gradient reversal, with a significantly larger fraction ultimately leaving the recording arena (Fig. 7D; Supplementary Fig. 7C). By contrast, the fraction of *eat-4* cKO animals leaving the arena did not differ significantly from wild type, despite their impaired progression toward the goal (Supplementary Fig. 7B– C). Thus, the enhanced negative thermotaxis of *unc-17* cKO animals does not reflect improved goal-directed navigation; rather, cholinergic transmission is required to keep navigation constrained around the learned goal instead of allowing continued movement beyond it.

This distinction is notable in light of recent work showing that AFD selectively amplifies temperature-change signals within a gain region centered on the learned thermal preference. This experience-dependent gain allows a derivative-based sensory code to remain referenced to an absolute behavioral goal (García et al., 2026). Our results suggest that co-transmitter control operates within this goal-centered sensory architecture: glutamatergic transmission promotes progression toward the goal, whereas cholinergic transmission restrains continued progression once animals reach the goal-associated region. Because these two transmitter systems can themselves be independently remodeled by sensory experience, their complementary behavioral functions provide a mechanism through which AFD could separately tune the drive toward a goal and the restraint required to remain near it. Such independent regulation may be especially important in changing environments, where the relationship between sensory input, position and behavioral goal must be continually updated. Adaptive co-transmission therefore offers a way to reweight distinct chemical outputs of the same sensory neuron while preserving both its sensory representation and underlying circuit connectivity, allowing behavior to remain referenced to a learned goal as the environment changes.

## DISCUSSION

Here we have shown that co-transmission provides a substrate for transmitter-specific synaptic plasticity within a single neuron. We arrived at this conclusion by following two endogenous small-molecule transmitter systems across multiple levels of organization—from their intracellular and subsynaptic distribution (Fig. 2), to their regulation by sensory experience, (Fig. 3 and 4) to their contribution to goal-directed behavior (Fig. 5-7). Glutamatergic and cholinergic machinery coexist within the same neuron and presynaptic boutons while retaining distinct organizational and regulatory properties, allowing sensory experience to differentially remodel the two transmitter systems rather than uniformly scaling the presynaptic terminal. Functionally, this partitioning allows the two transmitters to differentially regulate how a common sensory representation is translated into behavior, together supporting navigation toward and maintenance around a learned goal. We refer to this capacity to independently remodel coexisting transmitter systems in response to experience as adaptive co-transmission.

This transmitter-specific partitioning provides a cellular mechanism by which co-transmission can expand neuronal encoding capacity. Co-transmission has long been proposed to increase signaling flexibility through greater chemical diversity, and co-transmitters can differ in their release, postsynaptic actions and recruitment across contexts (Granger et al., 2017; Trudeau & El Mestikawy., 2018; Vaaga et al., 2014). What has remained less clear is whether such flexibility can arise from independent regulation of transmitter systems within the same presynaptic environment. Our findings show that it can: two coexisting transmitter systems can retain distinct organizational properties and be reweighted by experience. Encoding capacity may therefore increase not simply because a neuron possesses an additional chemical signal, but because each transmitter constitutes a partially independent dimension of synaptic output that can be differentially regulated according to sensory context. In this view, presynaptic partitioning creates the possibility for functional partitioning without requiring changes in circuit connectivity.

Acetylcholine–glutamate co-transmission in mammalian circuits also supports the idea that two transmitters from the same neuron can be differentially recruited. In habenular and striatal cholinergic systems, acetylcholine and glutamate make distinct contributions to synaptic signaling and behavior (Kljakic et al., 2022; Ren et al., 2011), indicating that co-transmission need not operate as a single, uniformly regulated output. Our finding that VGLUT/EAT-4 and VAChT/UNC-17 differ in their subsynaptic organization and dependence on conserved presynaptic machinery (Fig. 2) suggests that AFD could provide a genetically tractable platform for identifying mechanisms that selectively regulate each transmitter system. More broadly, these findings raise the possibility that examining co-transmitter systems across different sensory, physiological or behavioral contexts may reveal additional layers of transmitter-specific regulation across species.

Adaptive co-transmission also places co-transmission within a broader framework of neurotransmitter plasticity. Stable transmitter identity is actively maintained by transcriptional programs that sustain transmitter-specific gene expression in mature neurons (Deneris & Hobert, 2014; Destain et al., 2024; Serrano-Saiz et al., 2018), indicating that transmitter identity is actively maintained rather than irreversibly fixed. Neurotransmitter switching represents one manifestation of this flexibility, in which activity or experience can shift transmitter expression between functionally effective and ineffective levels, including coordinated gain and loss of transmitter programs within the same neuron (Pratelli & Spitzer, 2025; Spitzer, 2017). Our findings reveal a related but distinct form of plasticity: two transmitter systems can remain co-expressed while their relative contribution is independently reweighted by experience. Thus, adaptive co-transmission and neurotransmitter switching may represent complementary modes of transmitter plasticity through which neurons adjust chemical signaling without necessarily changing their underlying connectivity.

Context-dependent regulation of neurotransmitter usage across diverse systems suggests that this principle may extend beyond AFD. Acoustic experience induces dopamine-synthetic machinery in cholinergic auditory efferents, altering the potential balance of acetylcholine and dopamine signaling according to recent sound exposure (Wu et al., 2020), whereas photoperiod reversibly redistributes VGLUT3 across branches of serotonergic neurons to support adaptation to changing day length (Maddaloni et al., 2024). Acute stress can alter cotransmitter identity in serotonergic neurons, with consequences for generalized fear (Li et al., 2024). Recent work in glutamate/GABA co-transmitting inputs to the lateral habenula further shows that experience can change the relative functional contribution of two coexisting transmitters without eliminating either identity (Li et al., 2025). Together with our findings, these observations suggest that transmitter maintenance, spatial deployment and relative contribution represent multiple levels at which neurons can update chemical communication while preserving their underlying connectivity. The same flexibility can also become maladaptive: drugs of abuse can induce persistent changes in transmitter expression that contribute to altered behavior (Pratelli et al., 2024; Pratelli & Spitzer, 2025). Thus, mechanisms that normally allow transmitter systems to adapt communication to changing environmental demands may also create vulnerabilities when recruited by pathological experience.

Several mechanistic questions remain unresolved. Our data do not establish that VGLUT/EAT-4 and VAChT/UNC-17 occupy completely separate synaptic-vesicle populations, nor whether experience-dependent changes in transporter abundance translate proportionally into changes in transmitter release. The downstream mechanisms through which each transmitter alters individual navigation decisions also remain to be defined. Nevertheless, these limitations do not alter the central principle revealed here: two neurotransmitter systems sharing the same neuron can remain biologically distinct, can be differentially remodeled by sensory experience and can make complementary contributions to behavior across contexts.

Together, our findings establish adaptive co-transmission as a form of synaptic plasticity in which independently regulated co-transmitter systems reshape sensory information processing across contexts, expanding the functional versatility of neural circuits without changing their underlying connectivity.

## METHODS

### Strains

Worms were maintained at 20°C using standard techniques (Brenner, 1973) or placed at 15°C or 25°C when experimentally indicated. Strains were maintained on NGM plates seeded with *E. coli* (OP-50). The wild type (WT) is N2, and only hermaphrodite worms were used for this study. A complete list of strains appears below.

### Generation of new alleles

All allele and strain designs described below were generated in the Colón-Ramos laboratory by Andrea Cuentas-Condori, and genome engineering was performed by SunyBiotech.

CRISPR–Cas9 genome editing was used to insert conditional FLP-on tagging cassettes at endogenous vesicular neurotransmitter transporter loci. An mScarlet-I3 FLP-on cassette was inserted at the C terminus of *eat-4* to generate *eat-4(syb9207)*. Similarly, a SNAP-tag FLP-on cassette was inserted at the C terminus of *eat-4* to generate *eat-4(syb10661)*, and a HaloTag FLP-on cassette was inserted at the C terminus of *unc-17* to generate *unc-17(syb10698)*.

For AFD-specific recombination, SunyBiotech generated single-copy MosSCI transgenes expressing either FLP recombinase or Cre recombinase under the AFD-specific *gcy-8* promoter. These included *Pgcy-8::FLP* (*sybIs9557*; chromosome I, *ttTi4348*), *Pgcy-8::FLP* (*sybIs8656*; chromosome II, *ttTi5606*), and *Pgcy-8::Cre* (*sybIs9372*; chromosome I, *ttTi4348*).

Because *eat-4* and *tax-4* are separated by less than 0.2 cM, the *tax-4(p678)* null mutation was recreated directly in a strain carrying the *eat-4::GFP* FLP-on allele *syb8568*. The *tax-4(p678)* mutation converts the glutamine codon CAG at amino acid 82 to the stop codon TAG (Q82Stop). Introduction of the same CAG-to-TAG substitution into the *eat-4(syb8568)* background generated the *tax-4(syb9290)* allele.

### Confocal Microscopy

#### Imaging

Larval or young adult animals were immobilized on 10% agarose pads containing 10 mM levamisole. Live imaging was performed on a Nikon Ti2 microscope equipped with a CSU-W1 spinning-disk confocal head, an ORCA-Fusion BT sCMOS camera, a high-speed piezo Z-stage, and a 60×/1.40 NA Apo Lambda oil-immersion objective. Z-stacks were acquired at 0.3-µm intervals spanning the full depth of the AFD neuron closest to the coverslip. Imaging parameters, including laser power, exposure time, camera settings, and acquisition conditions, were kept constant across genotypes and experimental conditions within each imaging session. Animals cultivated at 20°C were included as wild-type controls in every imaging session and were used for session-specific normalization to account for day-to-day variability in fluorescence measurements.

#### Analysis

Images were analyzed using NIS-Elements AR software (version 6.10.01). Z-stacks were converted to maximum-intensity projections for quantitative analysis and two-dimensional rendering. Images were first subjected to rolling-ball background correction. Fluorescence thresholds for the VGLUT/EAT-4::GFP and VAChT/UNC-17::mRuby channels were established from the wild-type dataset and then applied identically to all images within the corresponding experiment.

Masks were generated independently for the EAT-4::GFP and UNC-17::mRuby signals, and the union of both masks was used to define synaptic regions of interest (ROIs). This common ROI mask was then applied to both fluorescence channels, allowing EAT-4 and UNC-17 fluorescence to be quantified from spatially matched synaptic regions.

For dual-labeled animals, GFP and mRuby fluorescence intensities were measured within each individual synaptic bouton along a single AFD axon. For each bouton, the UNC-17/EAT-4 fluorescence ratio was calculated, and bouton-level ratios were then averaged across all boutons within that axon to obtain one axon-level value per animal. Fluorescence measurements and fluorescence ratios were normalized to the corresponding mean value obtained from 20°C-grown wild-type animals imaged during the same experimental session. Normality was assessed using the Shapiro–Wilk test. For comparisons among three or more normally distributed groups, Brown– Forsythe and Welch ANOVA was followed by Dunnett’s T3 multiple-comparisons test. For comparisons between two groups, normally distributed data were analyzed using an unpaired Welch’s t-test, whereas non-normally distributed data were analyzed using the Mann–Whitney test.

### Live-imaging and analysis of synaptic protein trafficking

To simultaneously monitor trafficking of VGLUT/EAT-4 and VAChT/UNC-17 from the AFD soma into the axon, we used strain DCR10003 (*eat-4::mScarlet* FLP-on (*syb9207*) III; *unc-17::GFP* FLP-on (*ola503*) IV; *Pgcy-8::Flippase* (*sybIs8656*) II). Trafficking of endogenous RAB3/RAB-3 was analyzed using strain DCR9895 (*GFP::rab-3* FLP-on (*ox669*) II; *Pgcy-8::Flippase* (*sybIs8656*) I). Live imaging was adapted from Glomb et al., (2024)(Glomb et al., 2024).

Briefly, ten L4-stage animals cultivated at 20 °C were transferred to a 50-µl drop containing 0.5 mM levamisole and incubated for 20–40 min at 20 °C. Immobilized animals were mounted on 10% agarose pads containing 5 µl M9 buffer. Animals were screened using low-intensity illumination, and imaging was restricted to AFD neurons in which the process connecting the soma and axon was clearly visible and positioned closest to the coverslip.

Imaging was performed on a Nikon Ti2 microscope equipped with a CSU-W1 spinning-disk confocal head, two spatially aligned ORCA-Fusion BT sCMOS cameras for simultaneous two-channel acquisition, and a 60×/1.40 NA Apo Lambda oil-immersion objective. Single-plane images were acquired every 400 ms for at least 40 s using either 1 × 1 or 2 × 2 camera binning. Both binning conditions were used for VGLUT/EAT-4–VAChT/UNC-17 and RAB-3 recordings and did not measurably affect calculated transport velocities; measurements from both acquisition conditions were therefore combined.

Movies were analyzed in Fiji (Schindelin et al., 2012) (version 2.16.0). Image drift was corrected using StackReg, and kymographs were generated for each fluorescence channel using the MultiKymograph plugin. The line used to generate each kymograph was drawn along the AFD process with a width approximating the axon diameter and was consistently oriented from the soma toward the axon. Because the two fluorescence channels were acquired simultaneously using spatially aligned cameras, corresponding kymographs could be directly compared to assess whether EAT-4 and UNC-17 transport trajectories were coincident or independent.

Transport events were identified manually by an experimenter blinded to fluorescence channel. A continuous diagonal trajectory within a kymograph was scored as a single transport event. Spatial and temporal displacement of each trajectory were measured using the Straight Line tool and converted using the image pixel size and acquisition interval to calculate transport velocity (µm/second). Events moving from the soma toward the axon were classified as anterograde, whereas events moving toward the soma were classified as retrograde. For simultaneous EAT-4 and UNC-17 imaging, the reported event numbers represent the total number of transport events detected across all dual-color recordings. Transport velocity was calculated and analyzed for each individual event. Normality of event velocities was assessed using the Shapiro–Wilk test. For comparisons among three normally distributed groups, Brown–Forsythe and Welch ANOVA was followed by Dunnett’s T3 multiple-comparisons test. For comparisons among three non-normally distributed groups, a Kruskal–Wallis test was followed by Dunn’s multiple-comparisons test.

### STED live-imaging and analysis

#### Animal staining

For live STED imaging, we used animals carrying AFD-specific conditional tags for VGLUT/EAT-4, VAChT/UNC-17 and the active-zone protein Liprin-α/SYD-2: *eat-4::SNAP-tag* FLP-on (*syb10661*) III; *unc-17::Halo-tag* FLP-on (*syb10698*) IV; *syd-2::GFP* FLP-on (*wy1400*) X; *Pgcy-8::Flippase* (*sybIs8656*) II. Animals were labeled with Janelia Fluor dyes using a protocol adapted from Mueller et al., (2024)(Mueller et al., 2024).

To obtain synchronized animals, thirty-five L4-stage hermaphrodites were allowed to lay eggs from 10AM of day 1 to 6PM of day 2. After egg synchronization, adult animals were removed by lightly washing the plate with 1mL M9 and eggs remained on the plate with OP50 bacteria. Synchronized F1 eggs were allowed to develop for approximately 14-16 h at 20 °C to enrich for L2/L3 stage animals.

JF dyes were prepared as 1 mM stocks in DMSO. Briefly, 5 nmol of one dye was resuspended in 5 µl DMSO, and this suspension was then transferred to a dried 5-nmol aliquot of the second dye, generating a 5-µl combined stock containing 1 mM of each dye. For staining, L2/L3-stage animals were collected from plates using a glass pipette and transferred to 1.5-mL microcentrifuge tubes in 1 mL M9 buffer. Animals were pelleted by centrifugation at 3,000 rpm for 30 s and washed three times with M9 to remove residual bacteria. After the final wash, approximately 195 µl M9 + pelleted worms were retained, and 5 µl of the combined JF552 HaloTag ligand/JF669 SNAP-tag ligand stock was added, yielding a final concentration of 25 µM of each dye. Samples were protected from light with aluminum foil and incubated on a rotator at 20 °C for 2 h. Tubes were gently flicked every 15 min to minimize animal aggregation.

Following labeling, 1 mL M9 was added and animals were washed five times to remove unbound dye. After the final wash, approximately 200 µL M9 was retained, and stained animals were transferred using a glass pipette onto NGM plates seeded with OP50 bacteria. Animals were recovered at 20 °C and imaged 4–7 hours after staining.

#### STED imaging

Stained animals were immobilized on 10% agarose pads containing 10 mM levamisole. Live super-resolution imaging was performed on an Abberior MIRAVA Polyscope equipped with a 60×/1.4 NA oil-immersion objective. JF552-labeled VAChT/UNC-17 and JF669-labeled VGLUT/EAT-4 were imaged using 2D STED microscopy with a 775-nm depletion laser. The STED depletion pattern was aligned before each imaging session according to the manufacturer’s calibration procedure.

Endogenous Liprin-α/SYD-2::GFP fluorescence was used to identify presynaptic active-zone regions and define regions of interest (ROIs) along the AFD axon. STED images of the JF552 and JF669 channels were acquired simultaneously, followed by conventional confocal imaging of the GFP-labeled SYD-2 signal. Acquisition settings were kept constant across animals within each imaging session.

Raw 2D STED images were subjected to post-acquisition deconvolution using LightBox software (version w2510) with the manufacturer-recommended settings before quantitative analysis.

#### STED analysis

Deconvolved three-channel images were analyzed in Fiji (Schindelin et al., 2012). Images were first subjected to Gaussian smoothing using a sigma of 1. Fluorescence maxima were then identified independently in the Liprin-α/SYD-2::GFP, VGLUT/EAT-4::SNAP-JF669, and VAChT/UNC-17::Halo-JF552 channels using the Find Maxima tool.

For representative boutons shown in Figure 2A′, line scans were drawn from the active-zone SYD-2::GFP signal to the VAChT/UNC-17::Halo-JF552 signal, which represented the two most spatially separated fluorescence signals within the subsynaptic region. This orientation allowed visualization of the relative distributions of SYD-2, EAT-4, and UNC-17 across the bouton. For quantitative analysis, the fluorescence maximum was determined independently for each channel within individual synaptic boutons, and the distance between the maximum of each vesicular neurotransmitter transporter and the corresponding SYD-2 active-zone maximum was calculated for each bouton. Shapiro-Wilk test was used to determine samples were not normally distributed (p>0.05) and Mann-Whitney test was used for statistical comparison between both groups.

### Temperature training for confocal imaging

The day before, L4-stage animals were maintained overnight at 20 °C on food. The following day, plates were transferred to incubators set to the corresponding training temperature (15 °C, 20 °C, or 25 °C). A 10 mM levamisole solution used subsequently for immobilization was equilibrated in parallel at the corresponding training temperature. Animals were imaged after 1, 2, 3, or 4 h of temperature exposure, as indicated for each experiment (Fig. 3F and associated supplementary panels). For imaging, animals were immobilized with 10 mM levamisole on 10% agarose pads and imaged by confocal microscopy as described above.

### Thermotaxis behavioral assay

The day before each assay, L4-stage animals were maintained overnight at 20 °C on food (OP50 bacteria). The following day, 4–5 h before behavioral testing, plates were transferred to incubators set to the corresponding training temperature (15 °C, 20 °C, or 25 °C). Thermotaxis assays were performed as previously described(García et al., 2026; Hawk et al., 2018). A 10 × 10 cm agar assay pad was placed on a metal thermotaxis stage generating a linear 15–25 °C thermal gradient across the 10-cm arena and allowed to equilibrate for 5 min before animals were loaded. Temperatures across the arena were independently verified before each experiment using a high-accuracy thermometer (Fluke 54 II B) equipped with 80PK-1 bead thermocouple probes.

Animals were transferred to the center of the thermotaxis arena using one of two loading methods. In the first method, animals were transferred to an unseeded plate and allowed to crawl briefly before the plate was flooded with M9 buffer. Animals were then transferred onto the midline of the assay pad in three to four M9 droplets, with approximately three to four animals per droplet. In the second method, three 10-µL droplets of M9 were placed on bacteria-free regions of a plate, and animals were transferred sequentially through each droplet using a metal pick to remove residual bacteria. Three 4-µL droplets of M9 were then placed at the center of the thermotaxis arena, corresponding to approximately 20 °C, and three to four animals were transferred into each droplet using a metal pick.

Animals were monitored using a top-mounted digital camera (Mightex BCE-B050-U) while they exited the loading droplets, for no longer than 2 min. If most animals remained within the droplets after 2 min, excess liquid was gently wicked away using a clean Kimwipe. Image acquisition was then initiated immediately at 2 frames s^−1^ and continued for 30 min as animals navigated the thermal gradient. The first 15 min of each recording were used for behavioral analyses. Trajectories were analyzed using an adaptation of the MagatAnalyzer software package as previously described (Gershow et al., 2012; Hawk et al., 2018; Luo et al., 2014).

### Thermotaxis trajectory and occupancy analysis

Animal trajectories were recorded for 15 min at 2 frames/s on a 15–25°C linear thermal gradient, with X position converted to temperature using the calibration 900–1800 pixels = 15–25°C. Assay plate was treated as the independent biological replicate. To define the wild-type stabilization region, each animal’s mean X position during minutes 10–15 was calculated for trajectories tracked for at least 50% of this interval. Animal means were then averaged within each plate, and the mean and standard deviation of the wild-type plate distribution were used to define the ±1, ±2, and ±3 SD temperature landmarks. The same fixed landmarks were applied to all genotypes.

For time-resolved analyses, trajectories with valid X tracking for at least 50% of the assay and a valid position during the first 10 s were retained. For each plate and time point, landmark occupancy was calculated as the fraction of currently tracked animals positioned on the colder side of negative-SD landmarks or the warmer side of positive-SD landmarks. Genotype occupancy curves and heatmaps were generated by averaging plate-level values; heatmaps were sampled at 1-min intervals. Occupancy curves were integrated over the full 15-min assay to obtain a plate-level AUC. WT was compared with *eat-4* cKO and *unc-17* cKO using Wilcoxon rank-sum tests, with Holm correction across the two WT-referenced comparisons within each landmark.

Mean temperature trajectories were calculated first for each plate by averaging animal positions at each time point and then across independent plates; curves are shown as mean ± SEM. Genotype differences in mean position during minutes 10–15 were evaluated using a Kruskal–Wallis test. Temperature-position histograms were probability-normalized descriptive distributions of all valid positions within the indicated time windows and were not used for inferential statistics. Example trajectories show the first 10 animals with valid X–Y tracking for each genotype, selected in their original track order without selection based on navigational behavior.

### Trajectory preprocessing and spatial registration

Animal X–Y trajectories were extracted from cleaned tracking files and placed on a common assay timeline, with each assay plate retained as an independent replicate. Because the position of the metal thermal stage varied slightly between recordings, X coordinates were registered across experiments using the midpoint between the two fixed thermostats flanking the assay as a physical landmark. The thermostat midpoint was measured for each recording, and the entire X trajectory was translated so that this reference point corresponded to X = 1350 pixels (X_aligned = X_raw + [1350 − X_thermostat midpoint]). Y coordinates were unchanged, and no rotation, scaling, interpolation, or nonlinear transformation was applied. Tracks shorter than 300 frames were excluded; unobserved time points were retained as NaN.

### Behavioral quantification and statistical analysis

Animal trajectories were recorded for 15 min at 2 frames/s and processed at 1 Hz. Position along the thermal gradient was converted to temperature using the linear calibration 900–1800 pixels = 15–25°C. Assay plate was treated as the independent biological replicate. Thermal migration velocity was calculated from consecutive valid positions as dT/dt, with negative values indicating movement toward colder temperatures and positive values indicating movement toward warmer temperatures. Interval velocities were averaged within each track and 1°C temperature bin, and plate-level values were obtained by averaging across tracks. Genotype effects across temperature were tested using hierarchical linear mixed-effects models including genotype, temperature bin, and their interaction, with random intercepts for plate and track. Temperature bins were included only when at least four independent plates per genotype were represented.

Focused analyses were performed in the 19– 21°C and 17–19°C regions. Reorientation episodes were defined by clustering reversal calls occurring within 2 s, with the first call marking reorientation onset. Thermal experience preceding a reorientation was quantified as the mean dT/dt during the preceding 2 s. Reorientation rates were calculated for each plate and dT/dt bin as the number of clustered reorientation onsets divided by qualifying tracked animal-time; plate/bin observations required ≥10 s of exposure, and inferential analyses required ≥4 independent plates per genotype/bin. Reorientation counts were analyzed using Poisson generalized linear mixed-effects models with log-transformed exposure time as an offset and plate as a random effect; an observation-level random effect was included when overdispersion was detected.

Runs lasting ≥5 s with mean speed ≥0.04 mm/s were classified by heading as going down the gradient (|θ| < 64°), isothermal (64° < |θ| < 116°), or up the gradient (|θ| < 116°). True exit rates were defined as transitions from the respective incoming direction to either of the other directional states, normalized by tracked animal-time in that incoming state, and were analyzed using Poisson models with exposure offsets. Reorientation transition matrices were calculated within each plate as P (outgoing direction / incoming direction, reorientation) and then averaged equally across plates. Three-genotype analyses used omnibus genotype tests followed by planned pairwise contrasts; multiple comparisons were controlled using Holm correction, with P<0.05 considered significant.

### Gradient-switch thermotaxis analysis

Animals were tracked for 30 min at 2 frames/s on a linear thermal gradient. During the first 15 min, 900– 1800 pixels corresponded to 15–25°C; the gradient was then inverted over approximately 3 min, such that from 18–30 min the same physical coordinates corresponded to 23–17°C. Animal position was analyzed in the fixed physical X coordinate so that trajectories remained spatially continuous across the gradient inversion, with temperature used to relabel the same spatial axis before and after the switch.

For mean-position trajectories, animal X positions were first averaged within each independent assay plate at each time point and genotype means were calculated across plates and displayed as mean ± SEM. All valid instantaneous animal positions were pooled within consecutive 3-min intervals and displayed as probability-normalized histograms.

To determine the number of tracks that left the recording arena, individual tracks terminating within the terminal 1°C region of the arena were manually classified as animals that left the arena; all remaining animals were classified as stayed. Classification was performed by a scorer blinded to genotype. The proportions of animals classified as left or stayed were compared between wild type and each mutant genotype using chi-square tests.

## Code and data availability

All custom MATLAB code used for behavioral preprocessing, spatial alignment of thermotaxis recordings, trajectory analysis, statistical analysis, and generation of the data-derived panels in Figures 5–7 and Supplementary Figures 5–7 is publicly available at: https://github.com/cuentaa1/Cuentas-Condori-etal-2026. The repository includes codes for:

- extracting and aligning X–Y trajectories across assay plates
- defining wild-type spatial landmarks
- quantifying thermotaxis and reorientation behavior
- analyzing the gradient-switch experiments
- README files describing required inputs, analysis parameters, and outputs.

Repository also includes processed source data required to reproduce the quantitative analyses and figures, including X–Y trajectory matrices, plate identifiers, and derived source-data tables. Raw tracking files are available from the corresponding author upon reasonable request.

### List of strains

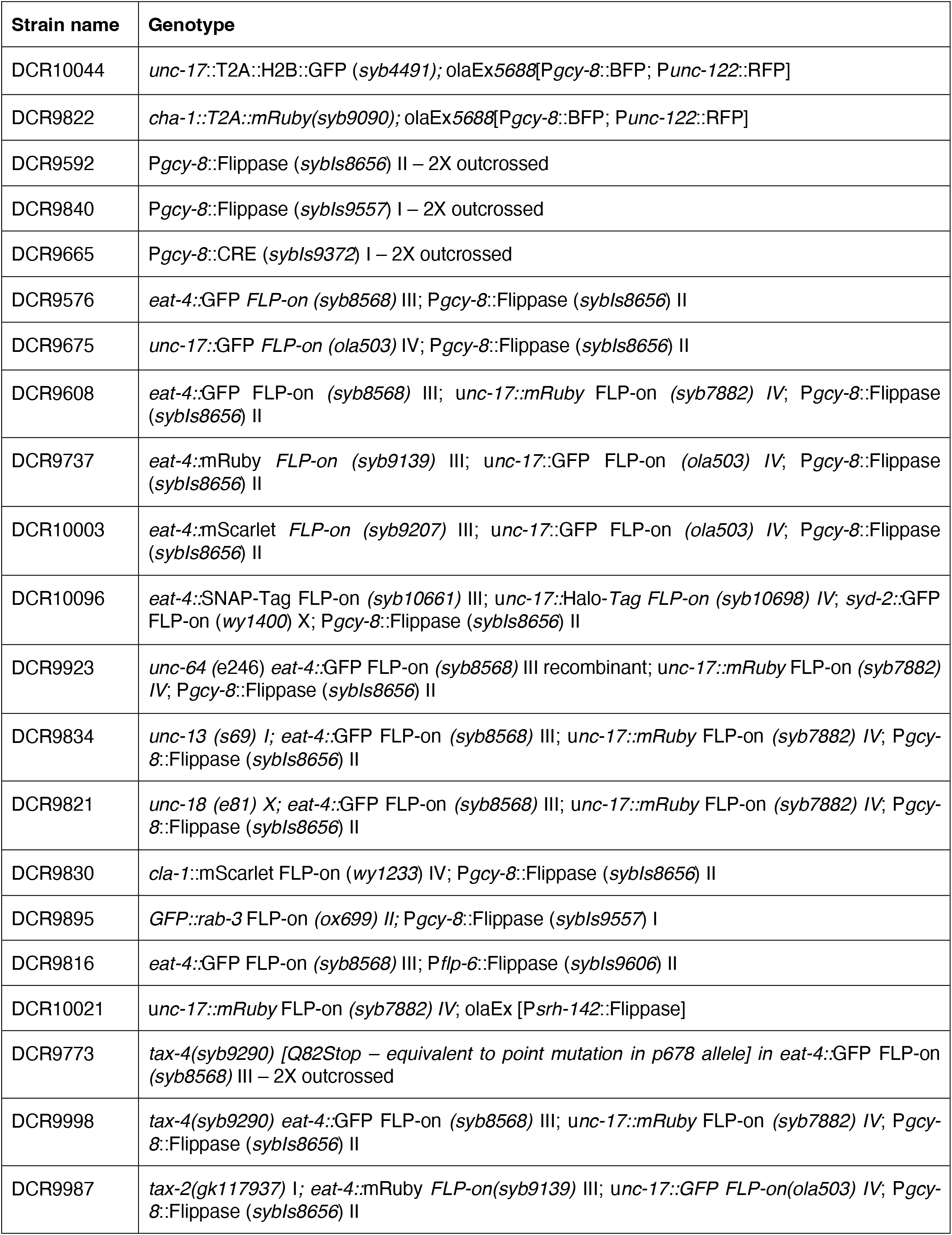
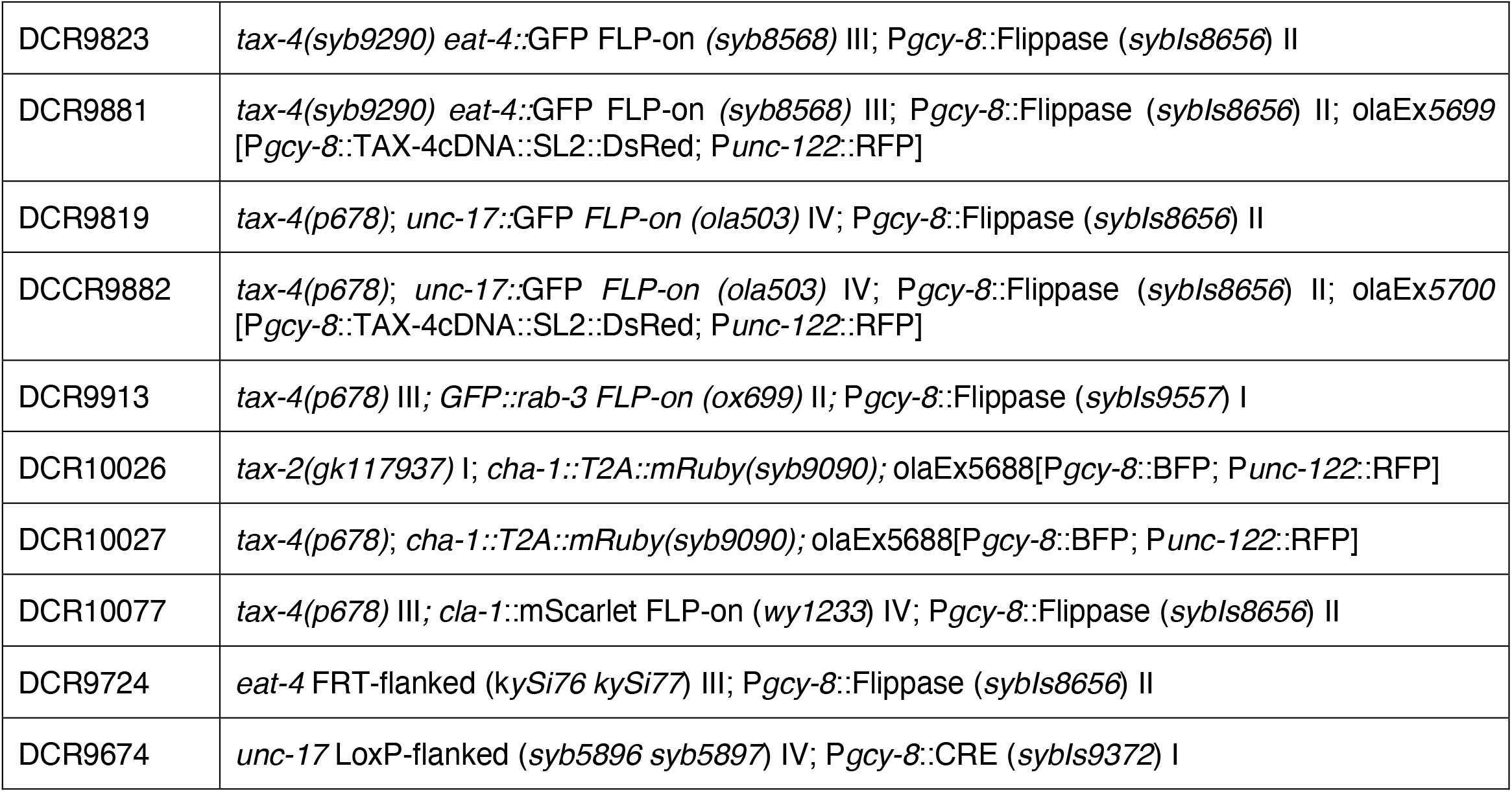

| Strain name | Genotype |
| --- | --- |
| DCR10044 | *unc-17::T2A::H2B::GFP (syb4491)*; *olaEx5688*[*Pgcy-8::BFP*; *Punc-122::RFP*] |
| DCR9822 | *cha-1::T2A::mRuby(syb9090)*; *olaEx5688*[*Pgcy-8::BFP*; *Punc-122::RFP*] |
| DCR9592 | *Pgcy-8::Flippase (sybIs8656)* II – 2X outcrossed |
| DCR9840 | *Pgcy-8::Flippase (sybIs9557)* I – 2X outcrossed |
| DCR9665 | *Pgcy-8::CRE (sybIs9372)* I – 2X outcrossed |
| DCR9576 | *eat-4::GFP FLP-on (syb8568)* III; *Pgcy-8::Flippase (sybIs8656)* II |
| DCR9675 | *unc-17::GFP FLP-on (ola503)* IV; *Pgcy-8::Flippase (sybIs8656)* II |
| DCR9608 | *eat-4::GFP FLP-on (syb8568)* III; *unc-17::mRuby FLP-on (syb7882)* IV; *Pgcy-8::Flippase (sybIs8656)* II |
| DCR9737 | *eat-4::mRuby FLP-on (syb9139)* III; *unc-17::GFP FLP-on (ola503)* IV; *Pgcy-8::Flippase (sybIs8656)* II |
| DCR10003 | *eat-4::mScarlet FLP-on (syb9207)* III; *unc-17::GFP FLP-on (ola503)* IV; *Pgcy-8::Flippase (sybIs8656)* II |
| DCR10096 | *eat-4::SNAP-Tag FLP-on (syb10661)* III; *unc-17::Halo-Tag FLP-on (syb10698)* IV; *syd-2::GFP FLP-on (wy1400)* X; *Pgcy-8::Flippase (sybIs8656)* II |
| DCR9923 | *unc-64 (e246)* *eat-4::GFP FLP-on (syb8568)* III recombinant; *unc-17::mRuby FLP-on (syb7882)* IV; *Pgcy-8::Flippase (sybIs8656)* II |
| DCR9834 | *unc-13 (s69)* I; *eat-4::GFP FLP-on (syb8568)* III; *unc-17::mRuby FLP-on (syb7882)* IV; *Pgcy-8::Flippase (sybIs8656)* II |
| DCR9821 | *unc-18 (e81)* X; *eat-4::GFP FLP-on (syb8568)* III; *unc-17::mRuby FLP-on (syb7882)* IV; *Pgcy-8::Flippase (sybIs8656)* II |
| DCR9830 | *cla-1::mScarlet FLP-on (wy1233)* IV; *Pgcy-8::Flippase (sybIs8656)* II |
| DCR9895 | *GFP::rab-3 FLP-on (ox699)* II; *Pgcy-8::Flippase (sybIs9557)* I |
| DCR9816 | *eat-4::GFP FLP-on (syb8568)* III; *Pflp-6::Flippase (sybIs9606)* II |
| DCR10021 | *unc-17::mRuby FLP-on (syb7882)* IV; *olaEx* [*Psrh-142::Flippase*] |
| DCR9773 | *tax-4(syb9290)* [*Q82Stop – equivalent to point mutation in p678 allele*] in *eat-4::GFP FLP-on (syb8568)* III – 2X outcrossed |
| DCR9998 | *tax-4(syb9290)* *eat-4::GFP FLP-on (syb8568)* III; *unc-17::mRuby FLP-on (syb7882)* IV; *Pgcy-8::Flippase (sybIs8656)* II |
| DCR9987 | *tax-2(gk117937)* I; *eat-4::mRuby FLP-on(syb9139)* III; *unc-17::GFP FLP-on(ola503)* IV; *Pgcy-8::Flippase (sybIs8656)* II |
| DCR9823 | *tax-4(syb9290) eat-4::GFP FLP-on (syb8568) III; P<sub>gcy-8</sub>::Flippase (sybIs8656) II* |
| DCR9881 | *tax-4(syb9290) eat-4::GFP FLP-on (syb8568) III; P<sub>gcy-8</sub>::Flippase (sybIs8656) II; olaEx5699 [P<sub>gcy-8</sub>::TAX-4cDNA::SL2::DsRed; P<sub>unc-122</sub>::RFP]* |
| DCR9819 | *tax-4(p678); unc-17::GFP FLP-on (ola503) IV; P<sub>gcy-8</sub>::Flippase (sybIs8656) II* |
| DCCR9882 | *tax-4(p678); unc-17::GFP FLP-on (ola503) IV; P<sub>gcy-8</sub>::Flippase (sybIs8656) II; olaEx5700 [P<sub>gcy-8</sub>::TAX-4cDNA::SL2::DsRed; P<sub>unc-122</sub>::RFP]* |
| DCR9913 | *tax-4(p678) III; GFP::rab-3 FLP-on (ox699) II; P<sub>gcy-8</sub>::Flippase (sybIs9557) I* |
| DCR10026 | *tax-2(gk117937) I; cha-1::T2A::mRuby(syb9090); olaEx5688[P<sub>gcy-8</sub>::BFP; P<sub>unc-122</sub>::RFP]* |
| DCR10027 | *tax-4(p678); cha-1::T2A::mRuby(syb9090); olaEx5688[P<sub>gcy-8</sub>::BFP; P<sub>unc-122</sub>::RFP]* |
| DCR10077 | *tax-4(p678) III; cla-1::mScarlet FLP-on (wy1233) IV; P<sub>gcy-8</sub>::Flippase (sybIs8656) II* |
| DCR9724 | *eat-4 FRT-flanked (kySi76 kySi77) III; P<sub>gcy-8</sub>::Flippase (sybIs8656) II* |
| DCR9674 | *unc-17 LoxP-flanked (syb5896 syb5897) IV; P<sub>gcy-8</sub>::CRE (sybIs9372) I* |

### List of plasmids

| Plasmid name | Genotype |
| --- | --- |
| DACR218 | *P<sub>unc-122</sub>::RFP* |
| DACR4202 | *P<sub>gcy-8</sub>::BFP* |
| DACR1876 | *P<sub>gcy-8</sub>::TAX-4cDNA::SL2::DsRed* |

## ACKNOWLEDGEMENTS

We thank Shaul Yogev for expert advice on the trafficking of synaptic proteins in intact animals; Hari Shroff and Mojtaba Tavakoli for discussions on imaging the subsynaptic organization of co-transmitter boutons; and Rachel Lee, Jesse Aaron, Teng-Leong Chew and Harikrushnan Balasubramanian at the Janelia Advanced Imaging Center for technical consultation and discussions on quantitative live imaging and image analysis. We thank Lin Shao and Mary Grace Velasco for training and expert advice on STED imaging and experimental design, and Joerg Bewersdorf for advice on quantitative analysis of super-resolution imaging. We thank members of the Yale StatLab for consultation on statistical analysis. We also thank members of the Yale Department of Neuroscience, the Yale Worm Club and members of the Colón-Ramos laboratory for helpful discussions and feedback throughout the project.

We thank Cornelia Bargmann (Rockefeller University), Steven Flavell (Massachusetts Institute of Technology), Oliver Hobert (Columbia University), Erik Jorgensen (University of Utah), and Kang Shen (Stanford University) for providing strains. Additional strains were provided by the Caenorhabditis Genetics Center (CGC), which is funded by the NIH Office of Research Infrastructure Programs (P40 OD010440). Figures were created in part using BioRender.com. STED imaging was performed using instrumentation in the Yale Wu Tsai Institute Microscopy Shared Facility (RRID: SCR_028114).

This work was supported by National Institutes of Health grants R35NS132156 and R01NS076558 to D.C.-R., and by a Pew Latin American Fellowship (AWD0006561), a Jane Coffin Childs Memorial Fund for Medical Research Fellowship (AWD0006564), and a Howard Hughes Medical Institute Hanna H. Gray Fellowship (GT15993) to A.C.-C.

---

**Supplement 1.**
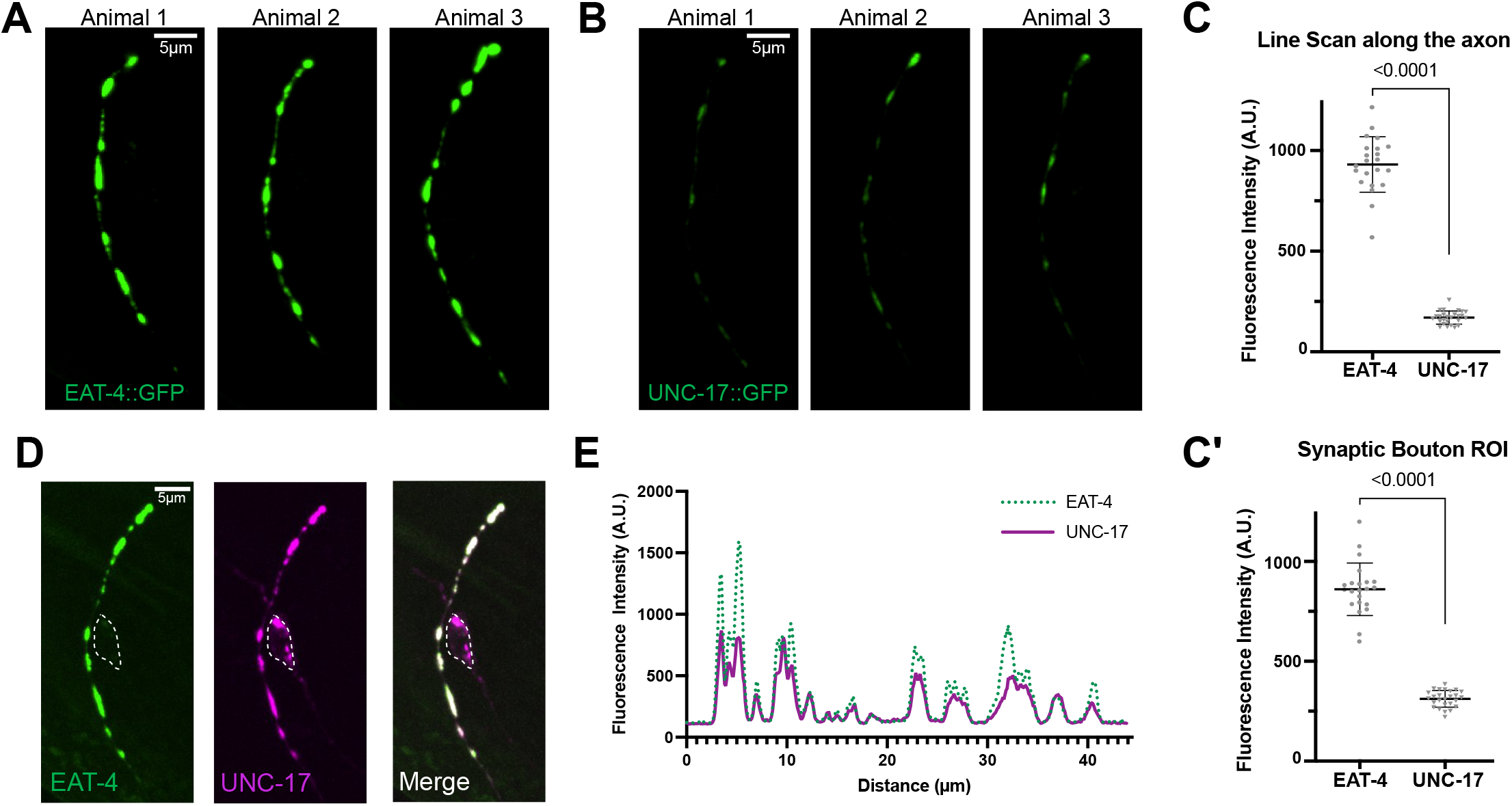
EAT-4/VGLUT and UNC-17/VAChT in AFD neurons. **(A)** Endogenous EAT-4::GFP in AFD axons of three independent animals. Scale bar is 5µm. **(B)** Endogenous UNC-17::GFP in AFD axons axons of three independent animals. Scale bar is 5µm. **(C)** EAT-4::GFP is more abundant than UNC-17::GFP when measures across AFD axons (p<0.0001) and **(C’)** in single AFD boutons (p<0.0001). **(D)** AFD axons localize EAT-4::GFP (green) and UNC-17::mRuby (magenta) to all vesicle clusters. Dashed structure labels the AFD soma. Maximum intensity projection. Scale bar is 5µm. **(E)** Line scan of EAT-4/VGLUT (green) and UNC-17/VAChT (magenta) along the AFD axon show an overlapping pattern.

**Supplement 2.**
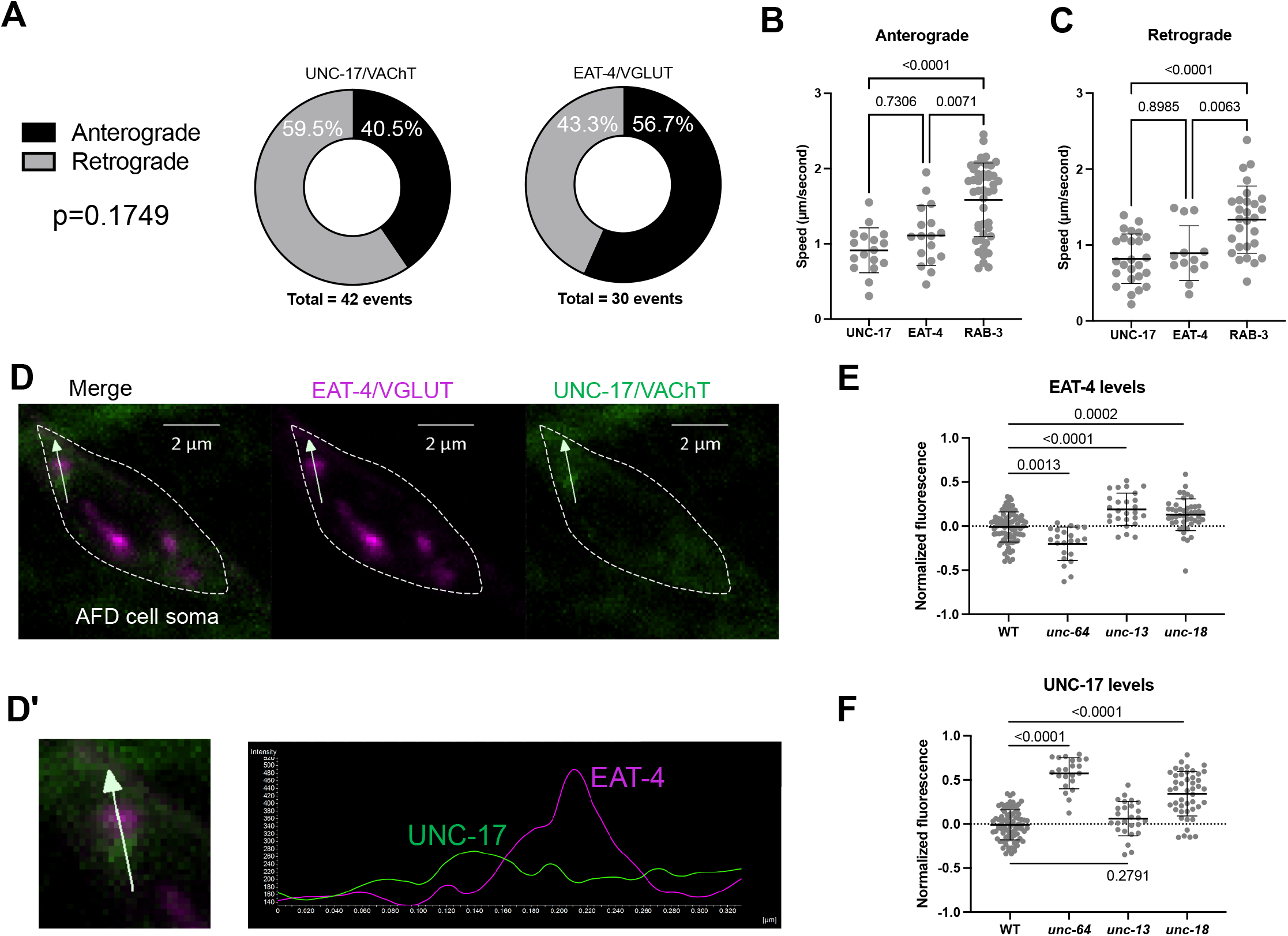
AFD neurons differentially organize and regulate EAT-4/VGLUT and UNC-17/VAChT. **(A)** Number of anterograde and retrograde events were not different between EAT-4/VGLUT and UNC-17/VAChT particles from the soma into the AFD axon (p = 0.1749). **(B)** Anterograde speed of UNC-17/VAChT is similar to that of EAT-4/VGLUT particles (p = 0.7306), but slower than RAB-3 particles (p < 0.0001). EAT-4/VGLUT particles also have slower anterograde particles than RAB-3 (p = 0.0071). **(C)** Retrograde speed of UNC-17/VAChT is similar to that of EAT-4/VGLUT particles (p = 0.8985), but slower than RAB-3 particles (p < 0.0001). EAT-4/VGLUT particles also have slower anterograde particles than RAB-3 (p = 0.0063). **(D)** EAT-4/VGLUT (magenta) and UNC-17/VAChT (green) in the AFD cell soma (dashed). **(D’)** Inset and line scan (arrow) on neighboring but non-overlapping EAT-4/VGLUT and UNC-17/VAChT in the AFD soma. **(E)** EAT-4/VGLUT is differentially regulated by synaptic proteins. Synaptotagmin/*unc-64* mutants have reduced EAT-4/VGLUT levels (p = 0.0013), while Munc13/*unc-13* (p<0.0001) and Munc18/*unc-18* mutants (p = 0.0002) accumulate EAT-4/VGLUT. **(F)** UNC-17/VAChT is differentially regulated by synaptic proteins. Synaptotagmin/*unc-64* (p<0.0001) and Munc18/*unc-18* mutants (p<0.0001) accumulate UNC-17/VAChT, while Munc13/*unc-13* mutants (p = 0.2791) maintain normal levels of UNC-17/VAChT at synapses.

**Supplement 3.**
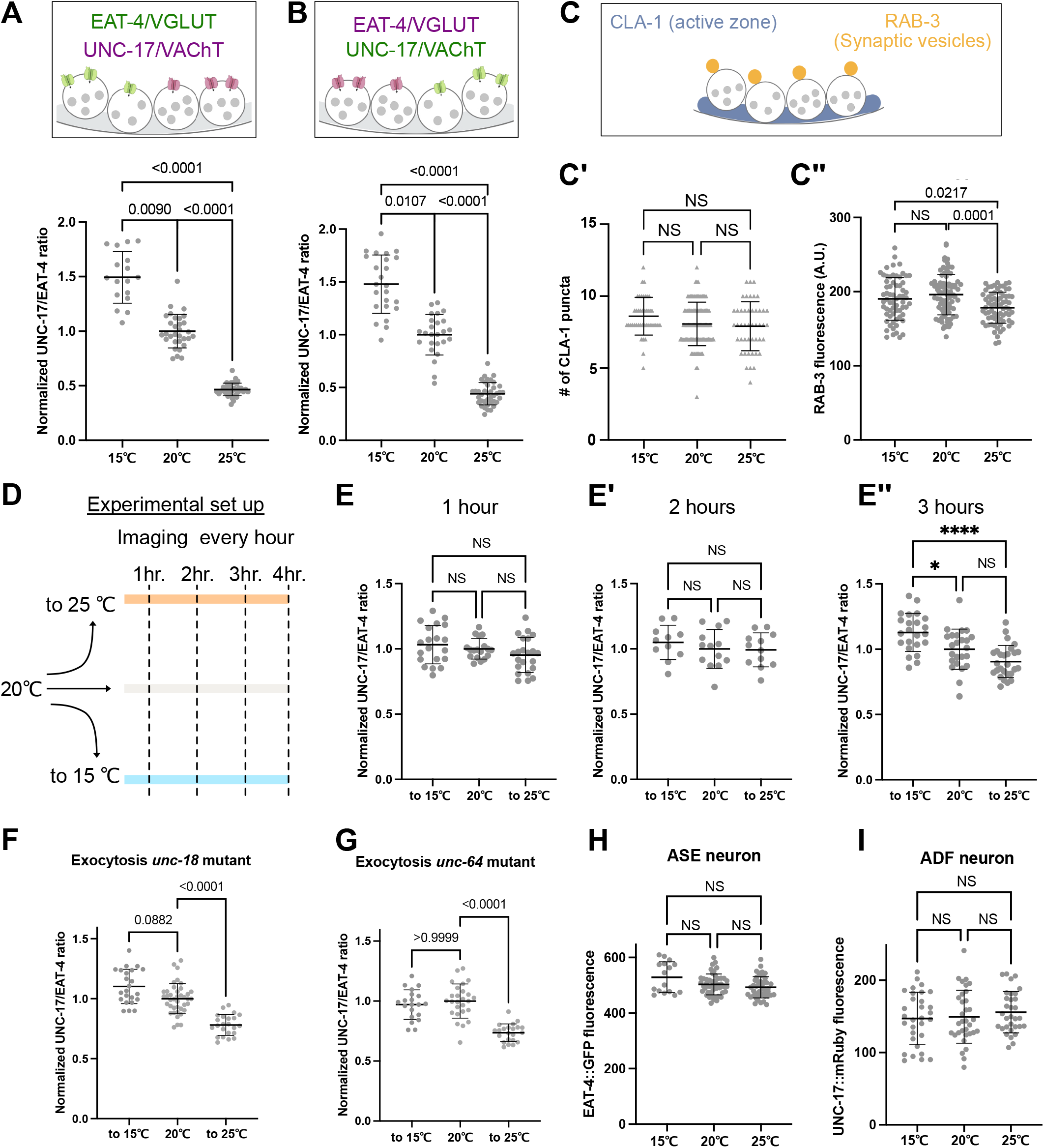
Experience-dependent regulation of AFD co-transmitter machinery is selective and time-dependent. **(A)** Quantification of AFD-specific UNC-17::mRuby/EAT-4::GFP ratios across three sensory experiences. **(B)** Quantification of AFD-specific UNC-17::GFP/EAT-4::mRuby ratio across three sensory experiences. **(C)** Quantification of AFD-specific (C’) CLA-1::GFP and (C’’) GFP::RAB-3 across three sensory experiences. **(D)** Experimental set-up to identify detection window of UNC-17/EAT-4 ratio changes. **(E)** Quantification of UNC-17::mRuby/EAT-4::GFP ratios after 1, (E’) 2 and (E’’) 3 hours of exposure to the new sensory experience. **(F)** Quantification of UNC-17::mRuby/EAT-4::GFP ratio after 4 hours of exposure in unc-18 mutants. **(G)** Quantification of UNC-17::mRuby/EAT-4::GFP ratio after 4 hours of exposure in unc-64 mutants. **(H)** Quantification of EAT-4::GFP fluorescence across three thermal experiences in the chemosensory neuron ASE. **(I)** Quantification of UNC-17::mRuby fluorescence across three thermal experiences in the chemosensory neuron ADF.

**Supplement 4.**
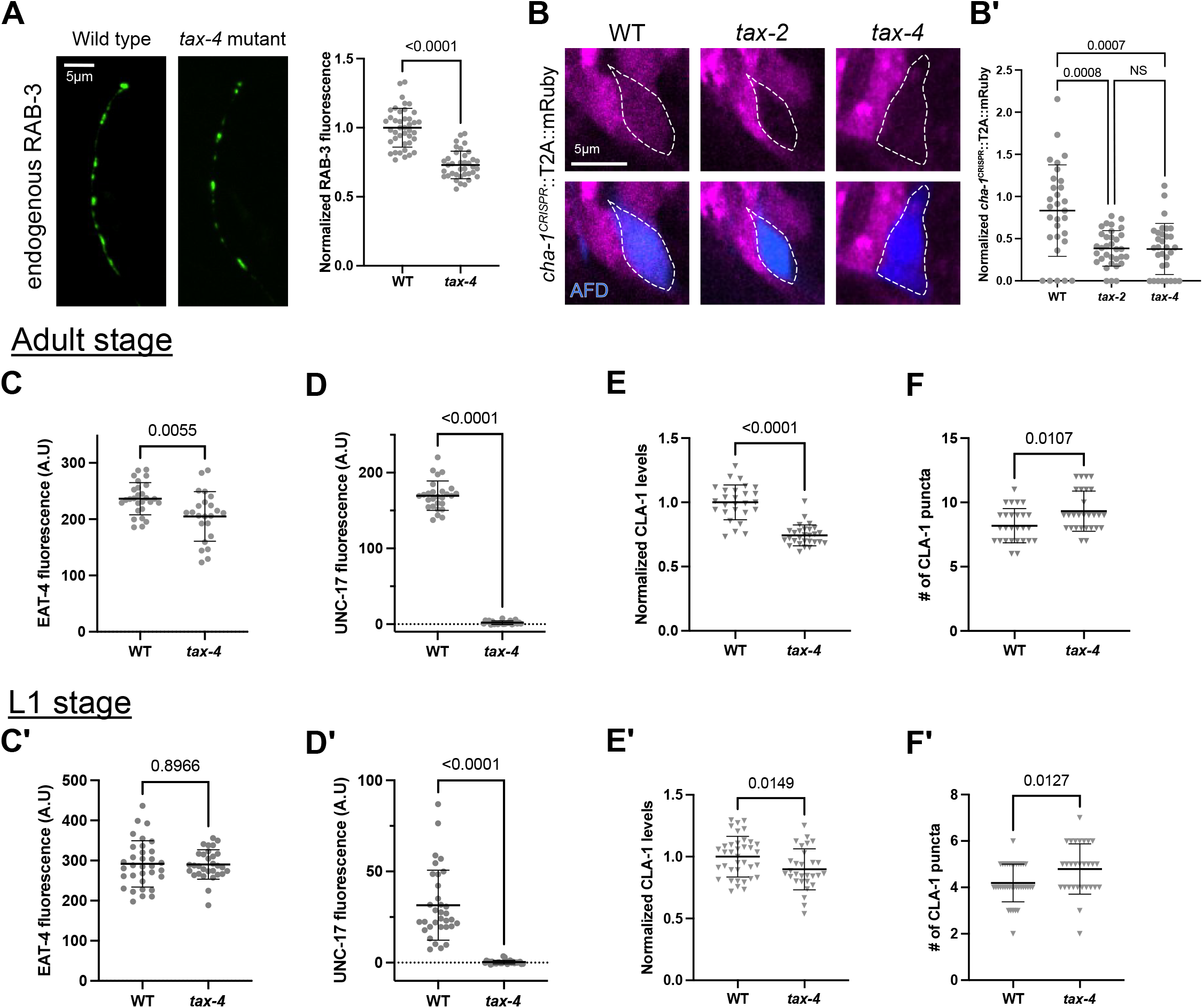
Sensory transduction regulates transmitter machinery and presynaptic organization across development. **(A)** AFD-specific GFP::RAB-3 levels in wild type and *tax-4* mutants. **(B)** Endogenous *cha-1* transcriptional reporter (magenta) in (left) wild type, (center) *tax-2* and (*right*) *tax-4* mutant animals. AFD neurons are labeled in blue. **(B’)** Quantification of cha-1::T2A::mRuby reporter in wild type, *tax-2* and *tax-4* mutant animals. **(C)** EAT-4/VGLUT fluorescence in wild type and *tax-4* mutant of **(C)** adult and **(C’)** L1 larvae. (D) UNC-17/VAChT fluorescence in wild type and *tax-4* mutant of **(D)** adult and **(D’)** L1 larvae. **(E)** CLA-1 fluorescence in wild type and *tax-4* mutant of **(E)** adult and **(E’)** L1 larvae. **(F)** Number of CLA-1 puncta in wild type and *tax-4* mutant of **(F)** adult and **(F’)** L1 larvae.

**Supplement 5.**
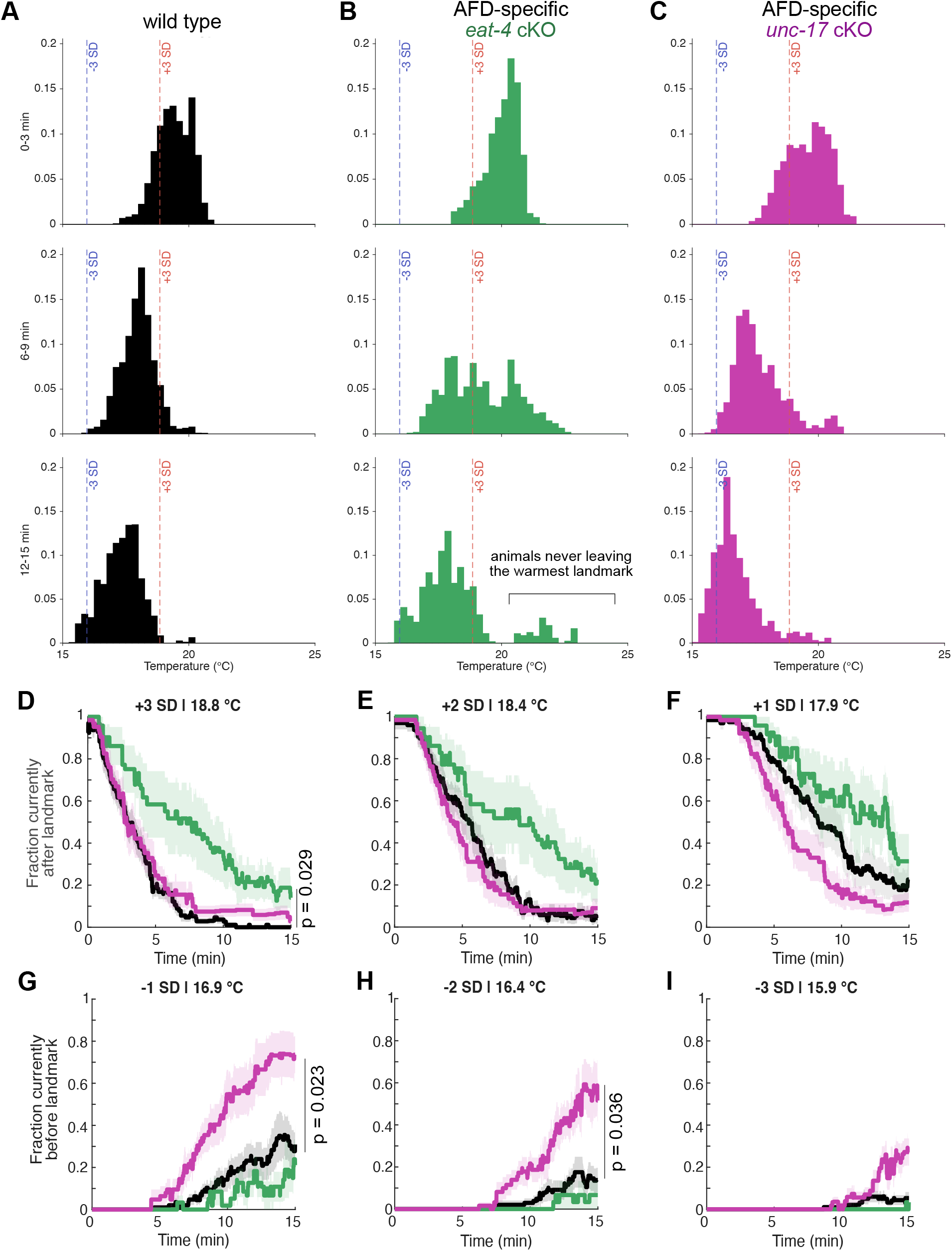
AFD glutamate and acetylcholine differentially shape progression across the thermal gradient. Histograms of X-positions of **(A)** wild-type, **(B)** AFD-specific *eat-4* cKO or **(C)** AFD-specific *unc-17* cKO mutant animals across three windows of time: (top) 0-3 minutes, (middle) 6-9 minutes, and (bottom) 12-15 minutes. **(D - I)** Fraction of wild type (black), AFD-specific *unc-17* cKO (magenta) and AFD-specific *eat-4* cKO (green) after **(D)** +3 SD, **(E)** +2 SD, **(F)** +1 SD, or before **(G)** -1 SD, **(H)** -2 SD or **(I)** -3 SD landmark.

**Supplement 6.**
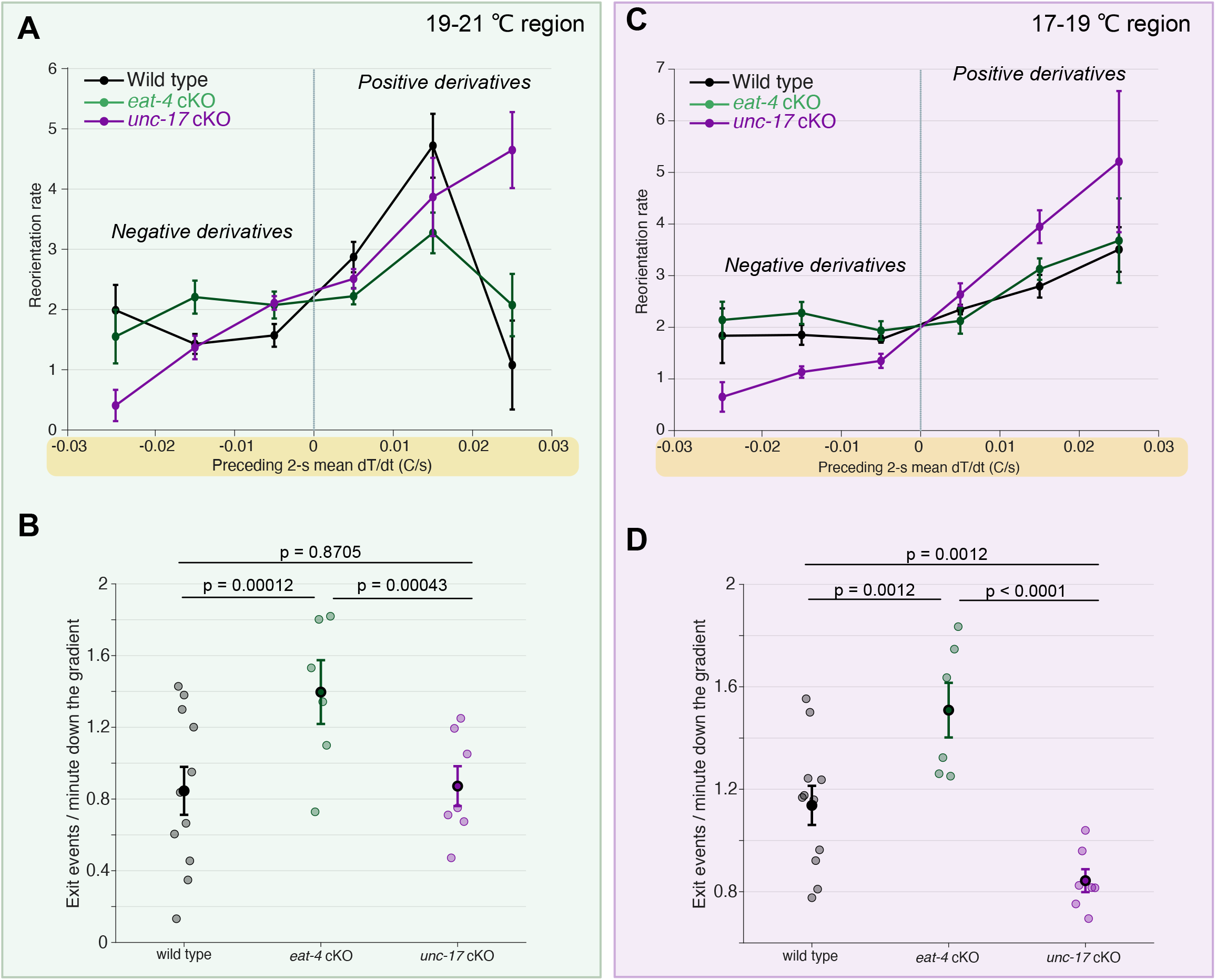
AFD glutamate and acetylcholine differentially regulate reorientation and run dynamics across thermal contexts. **(A)** Rates were analyzed using a three-genotype Poisson generalized linear mixed-effects model (GLMM) with genotype, dT/dt bin and their interaction as fixed effects, exposure time as a log offset, and plate as a random intercept. The genotype × dT/dt-bin interaction was evaluated by a joint Wald test (p < 0.0001). **(B)** Rate of exiting runs down the gradient within the 19–21 °C region for wild-type (black), AFD-specific *eat-4* cKO (green), and AFD-specific *unc-17* cKO (magenta) animals. Exit rates were analyzed using a three-genotype Poisson generalized linear model with genotype as the predictor and tracked time as a log exposure offset; each independent assay plate contributed one observation. Pairwise rate-ratio contrasts between genotypes were extracted from the same model and corrected across the three planned comparisons using the Holm method. *P* values shown are Holm-adjusted. **(C)** Reorientation rate as a function of the mean dT/dt experienced during the preceding 2 s for wild-type (black), AFD-specific *eat-4* cKO (green), and AFD-specific *unc-17* cKO (magenta) animals within the cooler 17–19 °C region. Statistical analysis was performed as in A. The genotype × dT/dt interaction was evaluated across all three genotypes (p < 0.0001). **(D)** Rate of exiting runs down the gradient within the 19–21 °C region for wild-type (black), AFD-specific *eat-4* cKO (green), and AFD-specific *unc-17* cKO (magenta) animals. Statistical analysis was performed as in B. *p* values shown are Holm-adjusted pairwise comparisons.

**Supplement 7.**
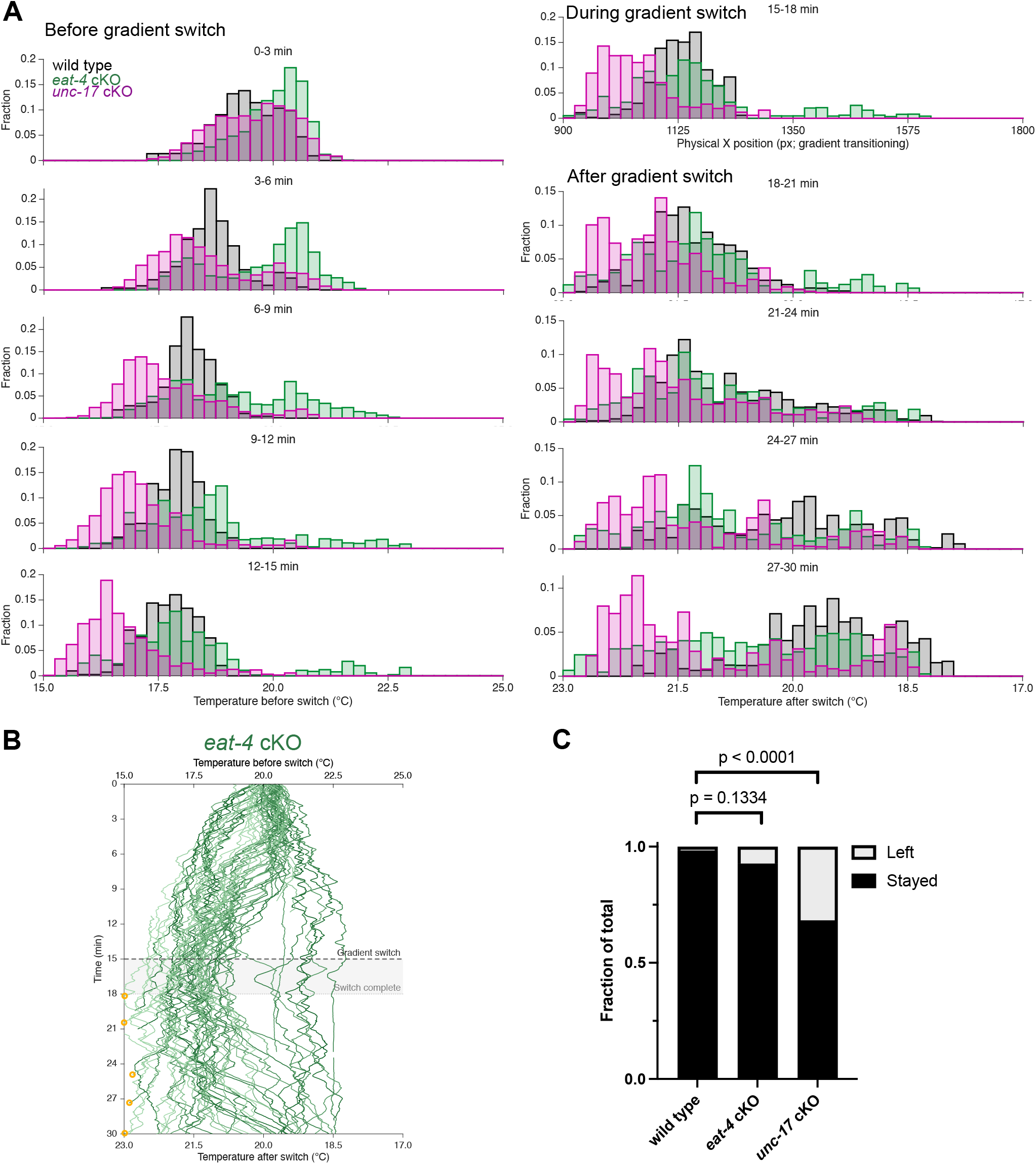
Population dynamics during adaptive tracking of a relocated thermal goal. **(A)** Distributions of instantaneous positions for wild-type (black), AFD-specific *eat-4* cKO (green), and AFD-specific *unc-17* cKO (magenta) animals in consecutive 3-min windows throughout the gradient-switch assay. For the 0–15-min period, positions are expressed as temperature on the initial 15–25 °C gradient. During gradient reversal (15–18 min), positions are plotted as physical x position because the thermal gradient was transitioning. After completion of the switch (18–30 min), positions are expressed as temperature on the reversed 23–17 °C gradient. Values represent the fraction of animals within each positional bin. **(B)** Individual thermal trajectories of AFD-specific *eat-4* cKO animals during the gradient-switch assay. Gradient reversal was initiated at 15 min and completed by 18 min. The upper x axis indicates temperature before the switch and the lower x axis indicates temperature after the switch. Orange circles indicate trajectories that exited the recording arena. *n* = 69 tracks from 6 independent assay plates. **(C)** Fraction of wild-type, AFD-specific *eat-4* cKO, and AFD-specific *unc-17* cKO trajectories that remained within or exited the recording arena during the assay. P calculated comparing two groups with a Chi-square test.

